# Systematic errors in connectivity inferred from activity in strongly coupled recurrent circuits

**DOI:** 10.1101/512053

**Authors:** Abhranil Das, Ila R. Fiete

## Abstract

Understanding the mechanisms of neural computation and learning will require knowledge of the underlying circuitry. Because it is slow, expensive, or often infeasible to directly measure the wiring diagrams of neural microcircuits, there has long been an interest in estimating them from neural recordings. We show that even sophisticated inference algorithms, applied to large volumes of data from every node in the circuit, are biased toward inferring connections between unconnected but strongly correlated neurons, a situation that is common in strongly recurrent circuits. This e ect, representing a failure to fully “explain away” non-existent connections when correlations are strong, occurs when there is a mismatch between the true network dynamics and the generative model assumed for inference, an inevitable situation when we model the real world. Thus, effective connectivity estimates should be treated with especial caution in strongly connected networks when attempting to infer the mechanistic basis of circuit activity. Finally, we show that activity states of networks injected with strong noise or grossly perturbed away from equilibrium may be a promising way to alleviate the problems of bias error.

## Introduction

Fully understanding the mechanisms of computation and plasticity in neural circuits requires knowledge of how neurons with specific functional properties are connected. Despite groundbreaking recent developments in direct circuit tracing^1–6^, obtaining connectivity data is difficult, expensive and slow. Hence the interest in developing and applying statistical methods to estimate connectivity directly^7–10^ from simultaneous activity recordings of many neurons within a circuit^11–13^.

A crude estimate of coupling strength between neurons can be made from correlations but, as is widely appreciated, correlations can arise from direct synaptic connections or a common input. Statistically sophisticated inference techniques – such as maximum entropy-based inverse Ising inference^8;14–17^, *l*_1_-regularized logistic regression^18^, and generalized linear models^7;19–22^ can ‘explain away’ correlations that arise from a common observed input, at least when all recurrently connected neurons are observed and the inference model exactly matches the model from which the data are drawn. This explaining away allows for deletion of a direct coupling that might have been drawn between a pair of neurons based on correlation alone (Fig. 1a), yielding connectivity estimates that are substantially sparser than the raw correlation graph.

These methods have been applied to data from low-level sensory circuits in the brain with excellent success in producing improved predictions of neural responses to stimuli, since they account not only for the influence of the stimulus but also for the collective influence of other cells in the network to neural activity^7;8;17;23;24^. Unfortunately, with the success of these models in activity prediction, it has been a common temptation, to which many have succumbed, to also interpret the inferred connectivity as biological connections^7;25–29^.

Consistently, the match between actual and inferred connections remains untested, even as the inferred connectivity continues to be referred to (by optimistic researchers) as a reasonable proxy for actual connectivity. There are two key requirements for the success of these methods in inferring connectivity, neither of which tend to hold in reality: all nodes must be observed, and the statistical model used to estimate connectivity must match very closely the model from which the data are generated. There is widespread recognition of the problem of unobserved nodes (frequently though it might be forgotten when interpreting inferred connections); therefore, we focus here on the second problem which arises from model mismatch. We show that the problem of model mismatch (and also partial observability of nodes) is large for recurrent circuits with strong weights.

We consider a natural question: When do statistically inferred weights in even fully observed circuits reflect true connections between neurons, and under what conditions can we expect the algorithms to (not) perform well in deducing biological connectivity? We hypothesize that in strongly recurrent systems, emergent phenomena – related to pattern formation and the presence of strong long-range correlations – present a fundamental challenge to the problem of explaining away in real data.

To study the question, we construct a simple structured neural network model with local recurrent connectivity whose strength can be dialed between weak (“sensory”) and strong (“memory”) regimes, and which in the latter regime produces emergent activity patterns with strong correlations between unconnected cells. The structured form of the connectivity is for convenience of easy visualization of patterns of error in the inference procedures. Structured connectivity makes it easy to visually distinguish between between activity correlations, inferred connectivity, and explaining-away errors relative to the generative ground-truth circuit. As we show later, results in the structured model generalize to unstructured (random) networks.

## Results

**Figure 1:**
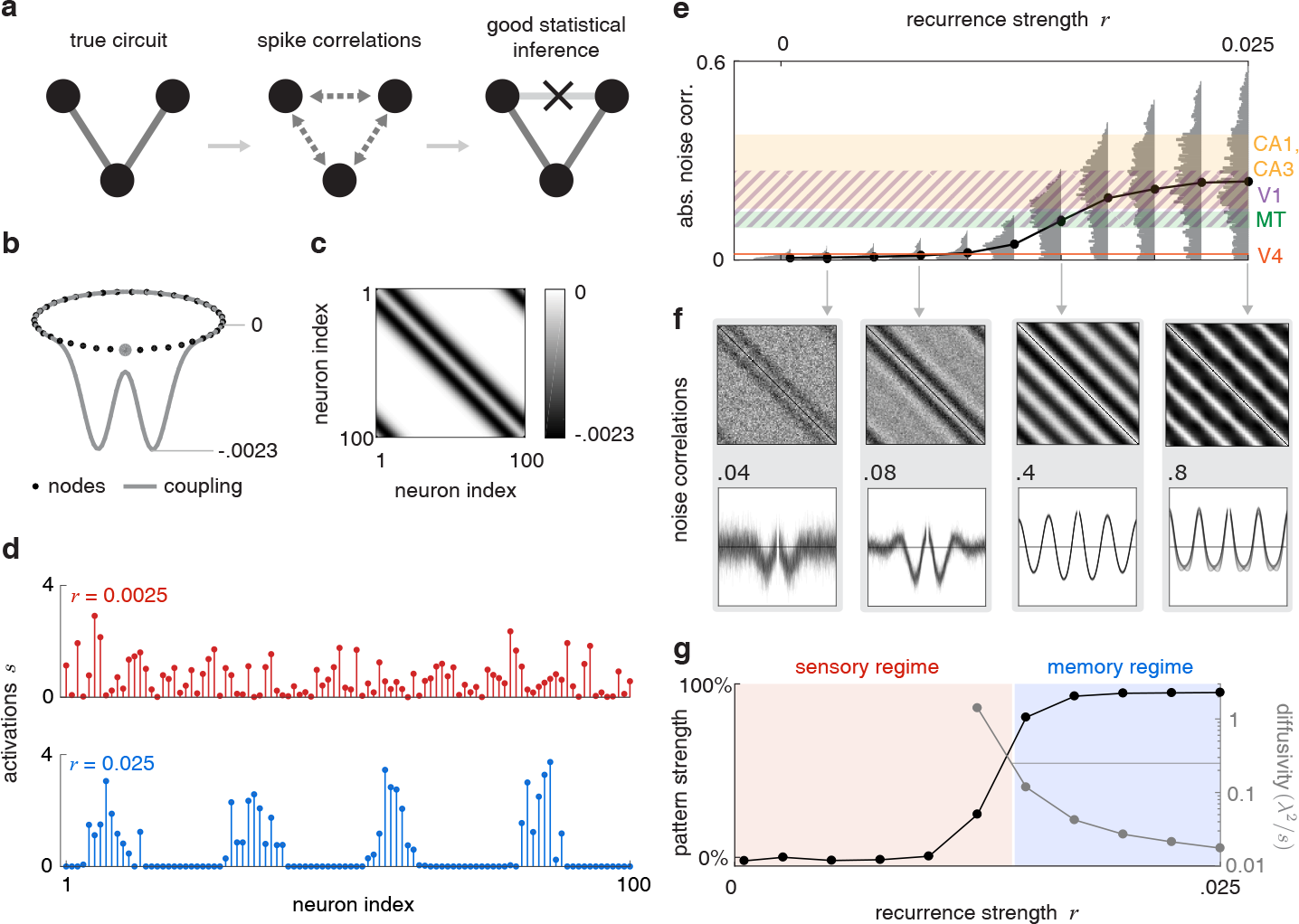
Structure and dynamics of the generative network. **a:** Left: A schematic three-neuron circuit with two excitatory connections. Center: All three neurons will be strongly correlated if the weights are large. Sophisticated circuit inference algorithms seek to ‘explain away’ the connection between the upper neurons by attributing their correlation to their common input instead of a direct connection. **b:** Neurons arranged in a ring-shaped network (black dots). Gray curve: Mexican hat- shaped weights from an example node (gray dot) to the rest. Each node in the network connects to the rest similarly. **c:** The resulting weight matrix **W** is circulant. **d:** Snapshots of synaptic activations of network cells with weak and strong weights. **e:** Absolute noise correlations between all pairs of neural spike trains binned at 10 ms (gray: histogram, black: means) increases with weight strength. For reference, horizontal color bands indicate experimentally measured values/ranges of noise correlation in several brain circuits. **f:** Top: Noise correlation matrix of the network cells at different weight strengths. Bottom: superposed vectors of noise correlation between each node and the rest of the ring. Numbers indicate the range of the vertical noise correlation axis on either side of 0 (horizontal line). **g:** Measures of the coherence and stability of neural activity pattern, against weight strength. Primary axis: strength of the periodicity of neural activity correlation. Secondary axis: diffusivity of the pattern phase at all weight strengths capable of sustaining a sufficiently strong pattern. Horizontal line marks 0.25*λ*^2^*/s*, corresponding to a state ‘memory’ of 0.5 s. We define the associated weight strength as the boundary between the ‘sensory’ and ‘memory’ regimes of the circuit.

### Generative network architecture and dynamics

Our generative model is a one-dimensional neural network of threshold-crossing spiking units interacting through rotation-invariant inhibitory recurrent connections **W** in a Mexican-hat profile on a ring (see Fig. 1b-c). Coupling is local: each cell inhibits nearby cells, but cells that are further away are unconnected. In addition to the recurrent input, all cells receive a uniform excitatory drive with a multiplicative noise component (see Methods). Cells generate spikes, which are used to drive the dynamics in the next time-step. The relative contribution of the recurrent inputs (relative to a background noise) can be dialed using a scalar weight strength *r* that multiplies the connectivity matrix **W**. When *r* is weak, network activity is noise-driven and uncorrelated, but increasing *r* transitions the network into a strongly interacting regime where it exhibits self-sustained periodic activity patterns. The pattern consists of multiple bumps, and unconnected nodes in different bumps exhibit strong long-range correlations (Fig. 1d).

In all that follows, inferred weights are compared with the true weight matrix after they are rescaled to obtain the best match (see Methods), thus changes in the size of inference error cannot be attributed to a simple mis-scaling of the inferred weights. Similarly, inference across weight strengths is performed on the same number of total spikes in networks with weak and strong weights, and inference differences cannot be attributed to different amounts of data.

To first quantitatively and qualitatively relate the weight strengths (*r*) to operating regimes of networks in the brain, we consider the effect of *r* on noise correlations, activity pattern coherence, and the stability of internal states over time.

Increasing the strength *r* of recurrent connections leads to stronger noise correlations (see Methods, Fig. 1e). These correlation values can be compared directly with measured values of the mean absolute pairwise noise correlation in various sensory and non-sensory brain areas including V1^30–32^, V4^33;34^, MT^35–38^, and hippocampus ^39^ (Fig. 1e, colored bands). As seen, medium to medium-large values in our range of *r* produce noise correlations consistent with primary and non-primary sensory processing stages in cortex; large values correspond to noise correlations from CA1 and CA3, areas associated with strong recurrent processing and memory. Medium-low and low values of *r* might correspond to weights that are weaker than found in even primary sensory areas because the correlations are smaller than seen in primary visual cortex. (But note that this comparison does not take into account the possibility of correlated noise inputs into primary sensory cortex^40–42^; thus we also use more qualitative measures, below, to relate values of *r* to different operating regimes in the brain.)

The structure of the noise correlation matrix, not just the size of the noise correlations, itself evolves with *r*: At weak *r*, where neurons are largely noise-driven, noise correlations are small and the noise matrix exhibits high variance with weak signatures of the true connectivity matrix (Fig. 1f, first panel). At intermediate *r*, the influence of noise decreases and activity correlations better reflect connectivity (1f, second panel). As *r* continues to increase, however, pattern formation sets in and noise correlations no longer reflect the true weights, capturing instead the correlations induced by the emergent periodic activity pattern (1f, last two panels).

Qualitatively, different values of *r* move the network along the sensory-memory continuum, defined by the coherence and diffusivity (see Methods) of activity states. Coherence is a measure of the fidelity of the shape of the activity profile over time (regardless of where the activity is centered); as *r* is increased, the network moves from a regime with zero pattern coherence to one with maximal coherence (Fig. 1g). The activity pattern itself drifts over time due to noise; this drift takes the form of a non-restorative random walk (OrnsteinUhlenbeck process), which we quantify by its diffusivity (see Methods). As weights increase in strength, the patterns become increasingly stationary and resistant to noise-induced drift, thus the diffusivity drops sharply (Fig. 1g). When diffusivity is low, the initial pattern phase is not rapidly lost and the phase can be used as a memory^43^. In Fig. 1g, the horizontal line marks the diffusivity value at which the RMS spread of the pattern phase equals half the pattern wavelength after 0.5 seconds; the starting phase is on average completely forgotten on this time-scale. The weight strength at this point is *r* = 0.0125; because both coherence and diffusivity change sharply around this value, we take it as our working boundary between the ‘sensory’ and the ‘memory’ regimes of the circuit. The sensory regime close to the memory boundary is highly-amplifying with slow responses.

### Inference at different weight strengths

We can now quantify the quality of circuit inference along the sensory-memory continuum defined above, beginning with the best-case experimental scenario, in which every neuron in the circuit is observable. We consider fully-observable data to highlight inference problems that arise in addition to the already well-known problems of inference in partially observed neural circuits.

Consider a dataset of spikes generated from the dynamical neural network model, with 10^8^ total spikes from the network at each value of *r* (see Methods). Because of its good performance in circuit activity prediction, at least at the sensory periphery ^7^, we apply a generalized linear model (GLM) to the data to extract an estimated weighted connectivity matrix, then measure inference error as the normalized *l*_2_-distance between the ground-truth and the properly-rescaled inferred weight vectors (see Methods).

When weights are weak, the recurrent connections have a small effect on neural activity relative to the ongoing noise, thus the signal-to-noise ratio (SNR) is low and the inferred connectivity exhibits uncorrelated errors (Fig. 2a, first panel), similar to the effects seen in the noise correlation matrix. The SNR improves, and inference error decreases with increasing weights (Fig. 2a, second panel), but only upto a point. Beyond this point, the inferred connectivity matrix begins to exhibit a new kind of error, visible as side-bands in the inferred matrix, last two panels of Fig. 2a (the distribution of inference errors, Supp. Info. §S.9 and Supp. Fig. S3 also shows a change in the pattern of errors). These errors are systematic (biased) overestimates of the existence of connections and magnitudes of weights between unconnected neurons. They result from a partial failure to explain away strong correlations, which are induced not by direct connections but by the emergence of stable activity patterns at the population level. Quantitatively, the GLM estimate is closer to the true connectivity matrix than is the raw noise correlation matrix, but the GLM estimate nevertheless exhibits a similar qualitative pattern of errors (compare Figs. 1f and 2a).

At any weight strength, the total inference error Δ can be decomposed into two orthogonal vector components: a variance Δ_*v*_ and a bias Δ_*b*_ (see Methods). As the weights increase in strength, the inference error vector rotates from being variance-dominated to bias-dominated (Fig. 2b). The total error is smallest for intermediate weights, when variance and bias contributions are both relatively low (Fig. 2c). The point of smallest error is far to the left of the sensory-memory boundary, in the very low weight regime (cf. regimes defined in Fig. 1e,g). When the inference model is not exactly matched to the data-generating model, as is typical, the minimal total error point is not an invariant quantity that reflects a critical point of the network dynamics or the inference process. Rather, the optimal inference point depends on the volume of data used for inference, as we show next.

### Variance but not bias errors decline with added data

The variance and bias errors behave differently as the volume of data available for inference is increased: examples at opposite ends of weight strength, where variance versus bias errors predominate, respectively, are shown in Fig. 3a, left versus right (total squared inference error is the sum of the squares of the variance and bias terms, 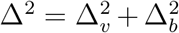). Data volume is defined as the total number of spikes used for circuit inference; because the average firing rate is held fixed, data volume is proportional to the time needed to collect the data. At the weakest weights, where error is nearly purely due to noise and thus variance-dominated, the error decreases inversely with data volume, as expected: 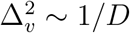, Fig. 3a inset at bottom-left with error well-fit by a line of slope −1 on a log-log plot (see Supp. Fig. S4a with fits and confidence intervals). Substantial bias errors arise for stronger weights, and these do not shrink with data volume, Fig. 3a, right (green area). Bias errors persist with data volume because they arise from highly-correlated neural activity states that are themselves highly structured and persistent.

It is clear from the full surface plots of total error (Fig. 3b), the relative fraction of bias errors (quantified by the normalized angle *θ*_*b*_ between the total error vector and the bias axis, Fig. 3c), and the bias and variance errors (Fig. 3d-e) that at all weight strengths the variance error decreases readily with added data while the bias term remains impervious. The bias error is therefore the asymptotic inference error in the limit of infinite data. Given that added data erodes the variance component of errors while bias errors are maintained, the point of minimum inference error should steadily move leftward with increasing data volume, as observed in 3f (dashed black line with arrow).

When using matched models for the data generation process and inference (e.g. using the Ising model for both), a scenario that is empirically implausible since we can never know the real model from which the data are generated, inference errors are simply variance errors at all weight strengths, and decay with data volume according to the same power-law (see Supp. Info. §S.1). In this case, the weight strength for best inference is an intrinsic critical point of the network (corresponding to maximal magnetic susceptibility in the Ising network, which in turn is directly related to the maximization of Fisher Information ^44^).

### Data required to perform accurate inference in memory networks is infeasible

From a practical or experimental perspective, the relevant question is how much activity data must be collected from a circuit to obtain an estimate of its connectivity at a specified precision, and how this value differs for sensory versus memory circuits.

**Figure 2:**
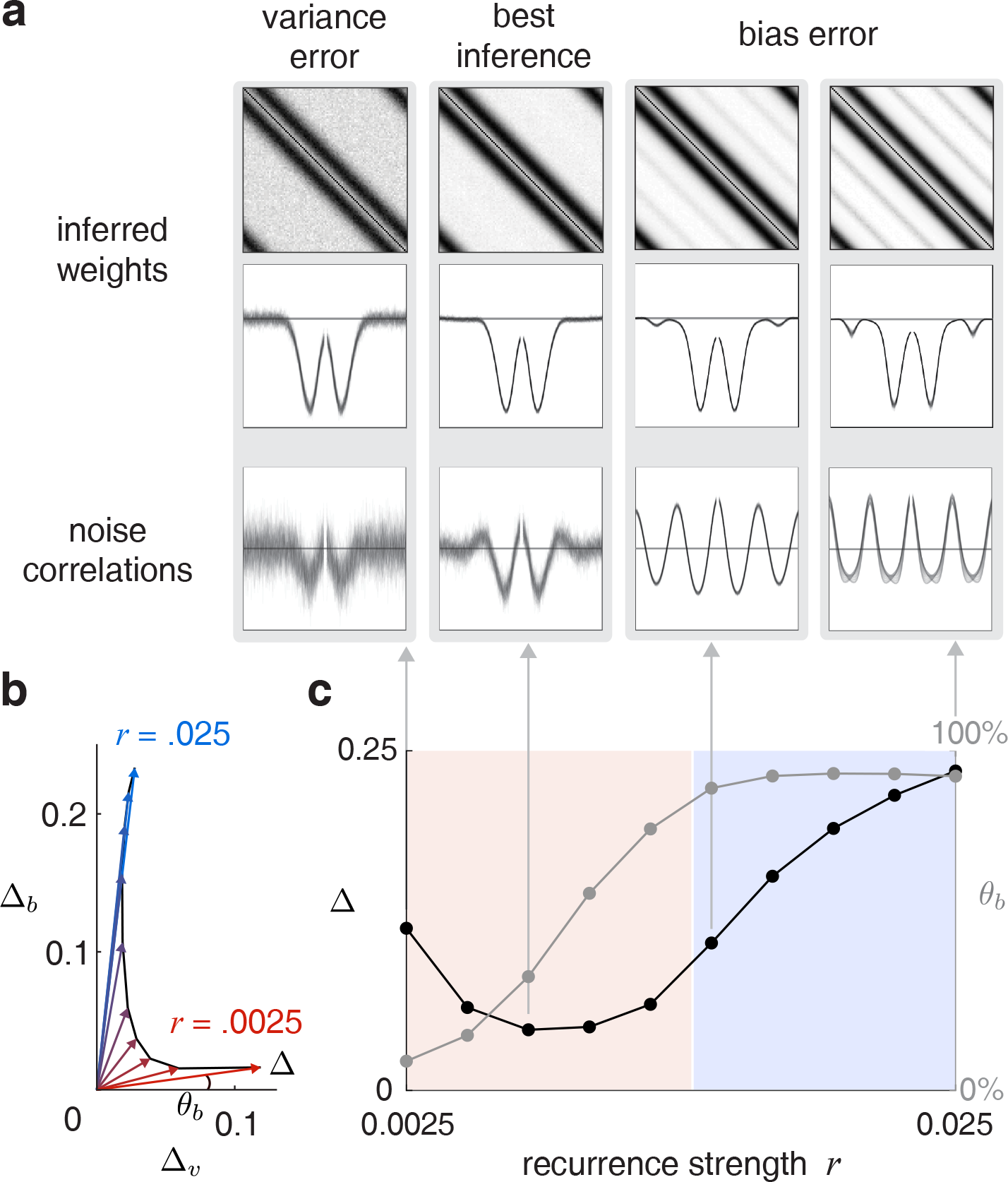
Circuit inference quality as a function of weight strength in a fully-observed circuit with 10^8^ total spikes (~ 6 hours of simulated spiking data at ~ 60 Hz avg. spiking rate per cell). **a:** Inferred weight matrices **Ŵ** (top row), superposed inferred weights from each node to the rest (middle row, line marks zero), and raw noise correlations (bottom row) for reference, at different weight strengths. **b:** Inference error (arrows) as a vector sum of orthogonal contributions from variance (Δ_*v*_) and bias (Δ_*b*_), for different weights. The magnitude of the vector is the total inference error Δ, and the normalized direction *θ*_*b*_ is the fraction of bias in the error. **c:** Total inference error Δ and fraction of bias *θ*_*b*_ against weight strength.

Thus, we fix the desired inference error (to 0.1194, corresponding to the inference error at *r* = .0025, horizontal line in Fig. 3f), and determine the data volume required, on average across multiple subsamplings of the data, to fall within 1% of this value. This process is equivalent to plotting a slice through the surface of Fig. 3b at a specified error level.

**Figure 3:**
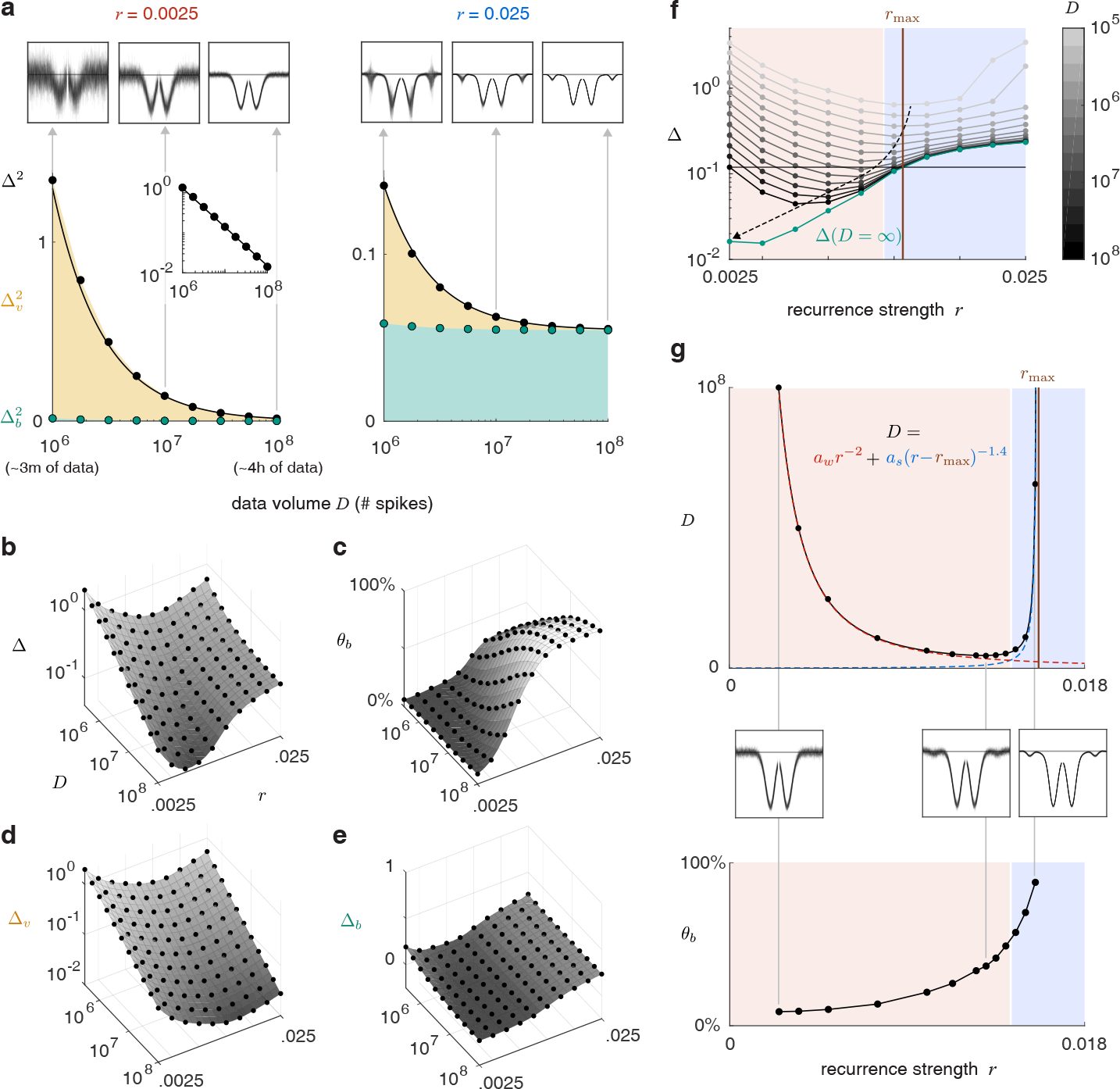
How data volume determines inference quality. **a:** Top: superposed weights from each node to the rest, inferred with different data volumes. Bottom: squared inference error Δ^2^, comprising squared variance error 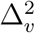 and squared bias error 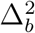, as a function of data volume (number of spikes collected). Black curve fits Δ^2^. Inset plot on log-log axes shows that for weak weights, Δ^2^ ~ 1*/D*. **b-e:** Full surfaces of total inference error, variance error, bias error, and bias error fraction, against weight strength and data volume. Black dots are data points, surface is an interpolation. **f:** Inference error against weight strength at different data volumes, and in the limit of infinite data (green). Optimal inference point moves towards *r* = 0 with increasing data (dashed arrow). **g:** Top: data volume required at different weights to meet a specified inference accuracy (horizontal line in plot f). Fitted black curve is a sum of two power laws diverging at *r* = 0 (dashed red) and *r* = *r*_max_ (dashed blue). Middle: superposed inferred connectivity from each node to the rest at different weight strengths, all corresponding to the same fixed inference error. Bottom: corresponding bias error fractions.

The data required to achieve a given inference error grows toward both extremes of weak and strong weights (Fig. 3g, black dots). As weights decrease, the influence of the recurrent connections shrinks and the data required to recover any information about the connections grows, diverging in the limit of vanishing weights. At the opposite end, the data demand escalates more steeply, and appears to diverge at a *finite* weight strength. We empirically fit this data demand curve with a sum of two power laws (3g, black, red, and blue curves). The first has a form 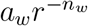, diverging at *r* = 0, while the other diverges at some finite weight *r*_max_ determined by the fit, and has the form 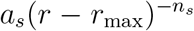. The best-fit yields the exponent *n*_*w*_ ≈ 2 for the first component, and for the second, *n*_*s*_ ≈ 1.4, diverging at *r*_max_ = 0.0157 (Fig. 3g, brown line). A network with stronger weights than *r*_max_ = 0.0157 cannot be inferred to the desired accuracy with *any* volume of data.

We can better understand the divergence of the data demand curve at a finite strong weight by plotting the obtained *r*_max_ on Fig. 3f: it corresponds to the point where the extrapolated infinite-data inference error (green curve) crosses past the criterion error (horizontal line), due to data-intractable bias errors at stronger weights.

### Results generalize across generative and inference models

In addition to using GLMs, we perform inverse inference on an Ising model (which corresponds to the maximum entropy model that fits the data means and covariances under the assumption of binary responses, see ^45^) using various techniques to obtain an estimate of connectivity from activity. Exact inverse Ising inference on a generalized Ising model (all-to-all connectivity with unequal weights) is NP-hard ^46^, requiring large amounts of data and intensive computation. Different algorithms fall at different points on a speed-accuracy tradeoff curve: Boltzmann machines perform optimal inference in the limit of large amounts of data if the data comes from the inference model class, but are slow. The simplest mean-field approaches are fast but suboptimal in accuracy.

We used an algorithm called minimum probability flow (MPF)^47^, which provides approximately optimal solutions at intermediate computational complexity and guarantees convergence to the correct (maximum likelihood) parameters in the asymptotic data limit.

Binarizing the data, as required for the method, results in information loss and poorer inference than GLMs and logistic regression (see Supp. Info. §S.3, Supp. Fig. S2). We performed inference at various bin sizes, and display results for the minimal-error result relative to the ground truth (Supp. Info. S.3, Supp. Fig. S2). Selection of the best result (and thus the best bin-size) cannot be done when the ground-truth is unknown, as is usually the case when doing circuit inference, but here it provides an upper-bound on the performance of such methods. (The best bin size, across weight strengths, is approximately the time-scale of single-neuron integration, suggesting that this is a good choice in general.)

At the other end of the speed-accuracy tradeoff for inverse Ising inference are mean-field methods, which are fast approximations. Among these we implemented the naíve mean field, the Thouless-Anderson-Palmer (TAP)^48^, and Sessak and Monasson (SM)^49;50^ approximations (see Supp. Info §S.4).

The naïve mean field method simply uses the negative inverse of the activity covariance matrix, so we consider it as the ‘raw correlations’-based connectivity estimate. This serves as a lower benchmark for circuit inference, and is plotted on Fig. 4a using spike count data as a comparison against the GLM and logistic regression (described next), and using binarized spike data as a comparison against the rest of the inverse Ising methods.

Next, we consider logistic regression and *l*_1_-regularized logistic regression^18;51;52^, in which the response of each neuron is regressed onto the activity of all the rest. The response variable must be binary, but the predictor values can have non-binary counts, thus we only binarize the spike data of the response neuron. Logistic regression achieves slightly better performance than the GLM, Fig. 4a.

**Figure 4:**
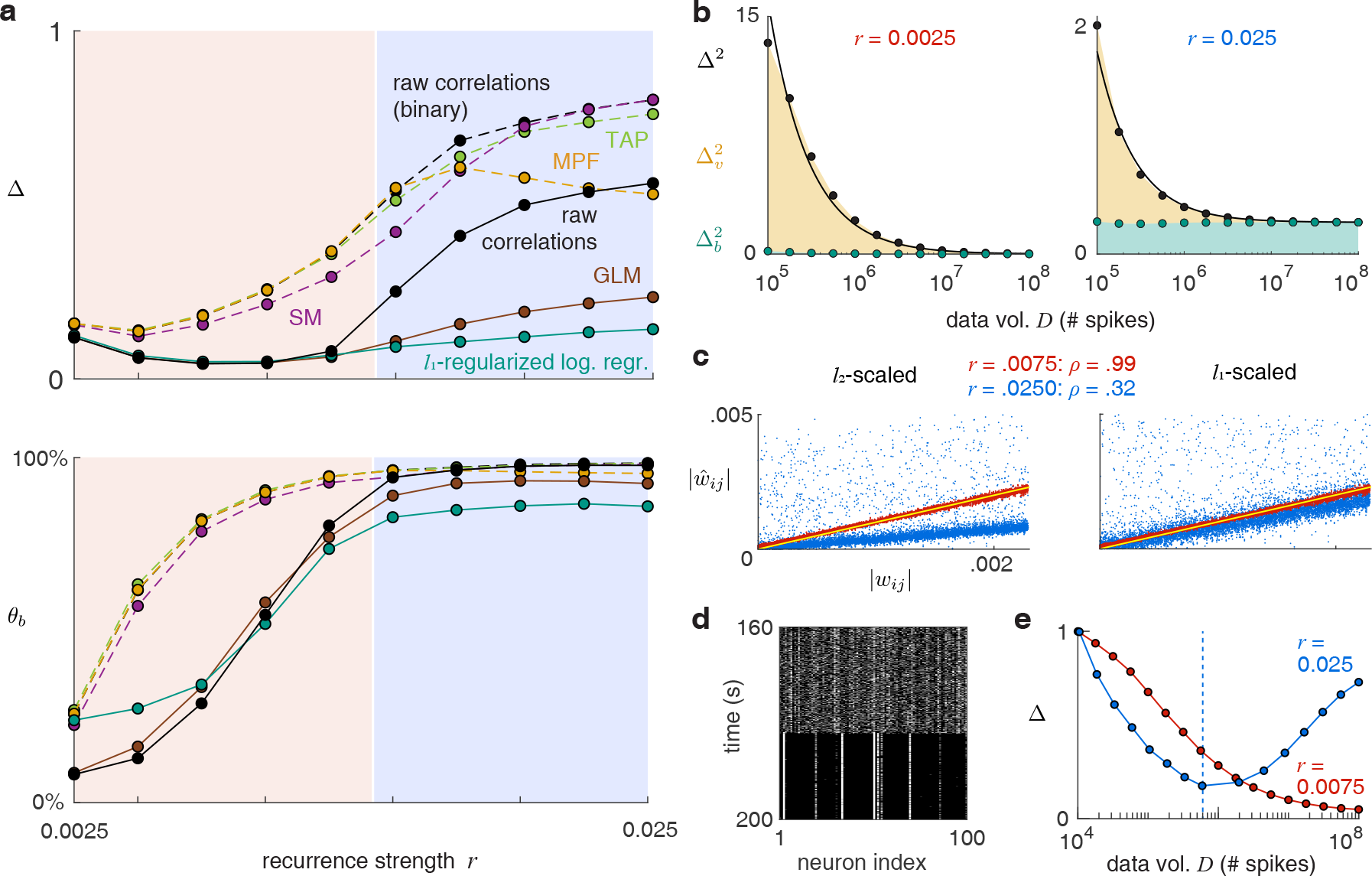
Results generalize across different inference methods. **a:** Results of inference on 10^8^ spikes from the network with different methods. Dashed lines are binary spike data-based inverse Ising methods. **b:** Inference error against data volume (similar to Fig. 3a) when using inverse Ising inference with minimum probability flow. **c:** Absolute inferred vs true weights and their correlation coefficients for a randomly connected network at weak and strong weights. Yellow line marks *y* = *x*. **d:** Spiking activity (white) in the random network with strong weights, initialized with weak activity, gradually evolves into a near-equilibrium state with a steady pattern (around 180s). **e:** Blue curve: For the evolving network in (d), we accumulate spike data over time, and perform inference as the data volume grows and the network gradually forms a pattern. Dashed line: onset of the steady activity pattern (near 180s) in (d). See Methods for more details. Red curve: the same process is repeated for a network with weak weights.

Since imposing an *l*_1_ penalty on the inferred weights prioritizes sparseness in the inferred weights^53^, and thus might push the network toward pruning or shrinking weights that make up the bias errors (the off-diagonal side-bands of Fig. 2a), we apply *l*_1_-regularized logistic regression^18^ (and tune the size of the regularization parameter at each weight to optimize inference accuracy, see Supp. Info. §S.8). However, *l*_1_-regularization does not eliminate side-bands and barely improves inference quality relative to unregularized regression (Supp. Fig. S5). Rather, *l*_1_-regularization forces all small weights to zero and retains larger weights, truncating or steepening the flanks of all peaks in the inferred weight matrix while retaining all peaks (including the side-bands).

When the recurrent weights are small, all inference methods, regardless of their statistical sophistication, perform equally well, and no better than the raw activity correlation matrix, at estimating connectivity. The main difference in the quality of the inference methods at low weights is the expected better performance of models that take spike counts rather than binarized spike trains into account, Figure 4a-b (solid versus dashed lines).

All methods replicate the qualitative trend of large inference errors at weak and strong weights, with best inference in the middle. At strong weights, all methods (with some mismatch) yield large bias errors that are impervious to additional data, Fig. 4b.

We switch generative models (while retaining the network architecture) then again perform inference, using a GLM with an exponential nonlinearity to fit data from both a linear-nonlinear-Poisson model and a GLM with rectifying nonlinearities. These produce the same qualitative patterns of error (see Supp. Fig. S6), showing that the results are not specific to a particular model for generating data.

Finally, the described effects are not specific to structured, low-dimensional ring-like circuit architectures, which we used to readily illustrate the pattern of bias errors. We consider next a network with all-to-all random symmetric connectivity. Connection strengths are drawn uniformly and randomly from the same range as the Mexican hat weight profile of the ring network. Circuit inference (here, using a GLM) displays the same patterns of errors as before, 4c (plots show the absolute values of actual and inferred weights, with inferred weights normalized according to two choices: minimizing the *l*_2_ or the *l*_1_ distances between the inferred and true weights).

Inference in the weak-weight regime is good, with inferred weights falling close to the unity line, but the strong-weight inferences deviate strongly from the true weights. Bias errors in connectivity estimation reside in the widespread scatter of inferred weights above the high-density blue region. These errors are reflected in the correlation coefficients of inferred vs true weights, which is a scale-independent measure of fidelity (*ρ* = 0.99 vs *ρ* = 0.32 for weak and strong weights, respectively, Fig. 4c).

In sum, the problem of overestimation of connectivity through bias errors at weight strengths that correspond to strongly amplifying sensory networks and to memory networks generalizes across models for data generation and inference, and across network connectivity architectures, whenever there is some mismatch in the inference model relative to the model generating the data (see Supp. Info. S.1 for inference in which the data generation model and the inference model are perfectly matched).

### Sampling data during non-equilibrium dynamics improves inference in strongly recurrent circuits

We consider what happens if activity data are sampled when the network is out-of-equilibrium. We initialize a network with strong weights (*r* = 0.025) far out of equilibrium, with low activity across neurons, Fig. 4d. The network states gradually evolve until they transition into an equilibrium activity pattern with strong correlations. As the states evolve, we collect activity data. Using the increasing amounts of data collected as the states evolve, but before pattern onset, improves inference, 4d. Interestingly, the slope with which inference improves with data volume is steeper in this non-equilibrium strong-weight network than for the in-equilibrium weak-weight network (due to higher SNR). Once the pattern emerges, however, the activity correlations contribute to bias error, and continuing to collect data in this regime now hurts inference, resulting in a worse match with the true weight matrix than if we used data collected only before pattern onset. This result shows that non-equilibrium activity data might result in much better inference results in strongly coupled recurrent neural networks ^54^.

### The circuit identification problem is inherent to activity states in strongly coupled circuits

The problem of circuit mis-identification when network weights are strong can be attributed to the very nature of the activity data in this strongly-coupled regime. Specifically, in the strong-weight regimes of strong sensory amplification or memory dynamics, the inherent distinguishability of spiking data generated from the true model and other candidate models is much smaller than when connections are weak. In other words, if different circuits generate convergently similar activity patterns, there is no inference method that would distinguish between them. Specifically, if the same neural network model generates data according to the true circuit and the mis-identified inferred circuit, how inherently similar or distinguishable are the resulting spike patterns?

Consider two weight matrices, **W**, our true circuit with only local synapses, and **W**′ with non-local synapses (Fig. 5). (**W**′ was obtained by inverse Ising inference on spike data generated by a dynamical neural network using the weight matrix **W** in the strongly recurrent regime, see Methods.) Our question is then: how inherently distinguishable are **W** and **W**′ in terms of the activity they produce at different recurrent weight strengths, independent of any inference procedure?

At each time-bin we record a vector ***σ*** of spike counts from all the nodes of a network, a spike word, which we may call the observed network state at that time. The distribution *p*(***σ***) of the states collected over time is the joint distribution of the activities of all the network nodes. The relative entropy (KL divergence) between the distributions produced by two circuits serves to measure the dissimilarity of the states they generate. This relative entropy measured between two parametrically interpolatable models is the Fisher Information contained in the data about the models (see Methods).

Such measures are accurate only when the individual distributions are well-sampled, and the state space of *N*-neuron networks is vast when *N* is even moderately large (here, *N* = 100). We employ two strategies to make the problem tractable: we consider only binarized spike data (Supp. Info. S.3), and instead of considering length-100 spike words, we consider spike words of smaller, 10-neuron segments of the network, whose lower-dimensional distribution we can sample sufficiently well (see Methods).

**Figure 5:**
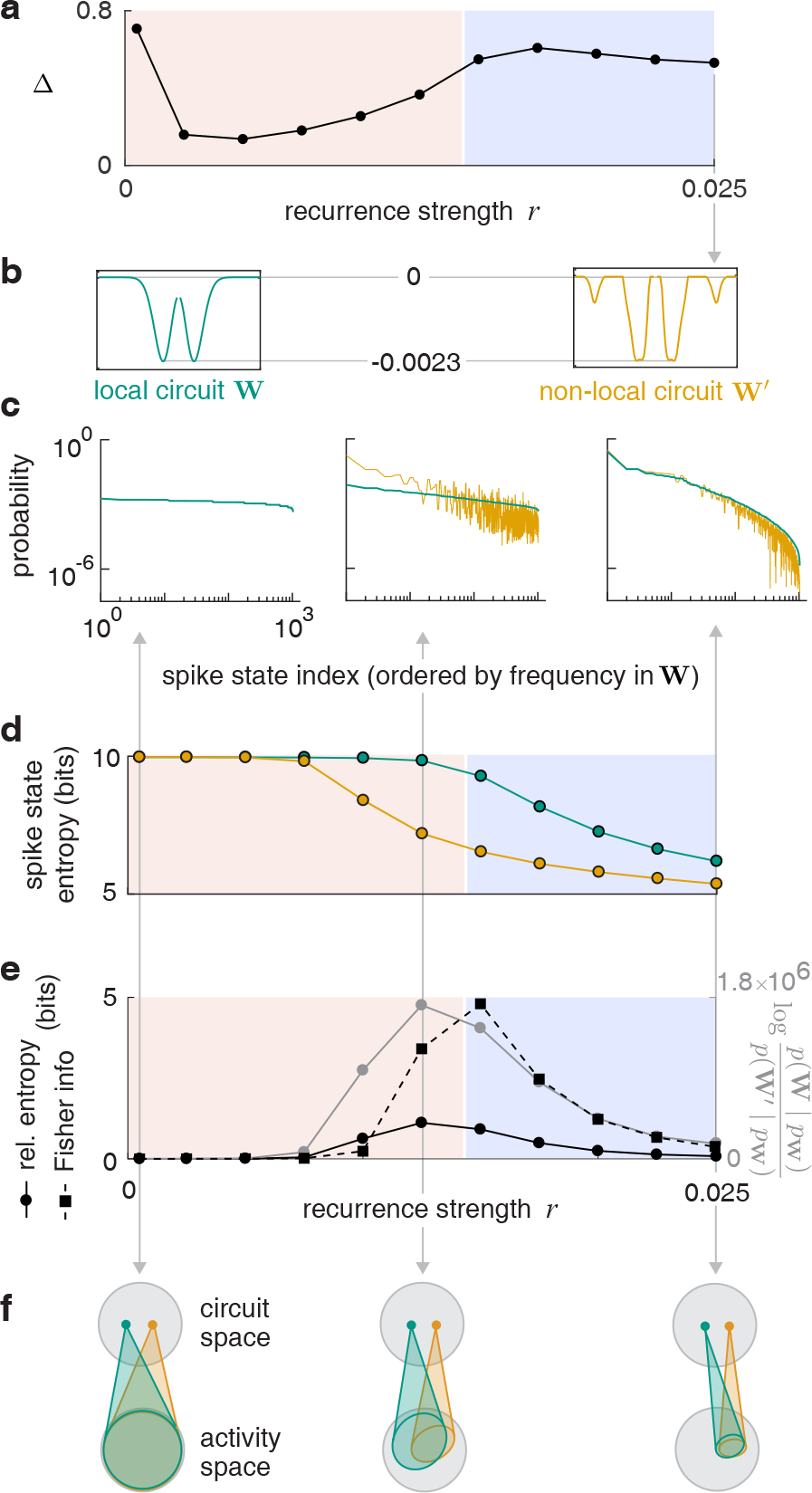
Recurrently stabilized activity patterns fundamentally limit circuit identification. **a:** Inference error using inverse Ising with MPF. **b:** True weight profile (with local synapses) of the generative circuit, and inferred weight profile (with non-local synapses) with strong weights (Fig. 2a). **c:** Frequency distributions of binarized spike state vectors of 10-neuron segments from the two circuits, ordered by their frequency in the local-circuit data, at different weight strengths. **d:** Entropies of the spike state distributions of the two circuits against weight strength. **e:** Relative entropy between the distributions, Fisher information, and the log likelihood ratio of the local vs non-local circuit given the local-circuit data, against weight strength. **f:** Schematic of mapping from circuit model space to activity data space, as weights vary.

The distributions (relative frequencies) *p***W**(***σ***) and *p***W**′(***σ***) of the 2^10^ different spike states ***σ*** of 10-neuron segments from the two circuits at different weight strengths are shown in Fig. 5c. The states are sorted along the horizontal axis according to their frequency in *p***W** (***σ***); the heights in green and yellow at a given point along the abcissa denote the relative frequencies of the same states ***σ*** in the activities of the two circuits.

When weights are weak, activity in both circuits is driven almost exclusively by the noisy feedforward input, thus state probabilities are very similar and flat in both models *p***W** and *p***W**′; the entropy of the states is the maximum of 10 bits (Fig. 5d). As recurrent strength is increased, both distributions contract in state space and their entropies drop. This occurs earlier in the **W**′ circuit because the added non-local recurrent couplings drive an earlier onset of strong correlations or patterning; thus *p***W**′ is more narrowly peaked (note logarithmic vertical scale) than *p***W** for weaker weights, and the two circuits are distinguishable. As the weights continue to increase, pattern formation catches up in **W**, while the state space of both models continues to shrink. However, the set of activity patterns and thus the state distributions converge again at strong weights, meaning that the two models become inherently less distinguishable, at least under the dynamics of the data-generating model. The same result holds when considering larger segments of the network (up to 22 nodes; see Methods and Supp. Fig. S7).

These distributional effects can be quantified with three closely related metrics, the relative entropy, the likelihood ratio of **W** vs **W**′ given the activity data *p***W**, and the Fisher information. (This measures the sensitivity with which one can determine the value of a continuous parameter in a generative model from the data. The parameter here is the linear interpolation variable *θ* in a generative circuit model where the recurrent connections are given by *θ***W** + (1 − *θ*)**W**′, see Methods). Larger values of these metrics indicate that data produced by the two circuits are inherently more distinguishable. It is clear from these measures that the alternative models are maximally distinguishable close to the sensory-memory transition, at the onset of patterning, and become more indistinguishable (for a fixed volume of data) at both stronger and weaker recurrent weights. This result is consistent with findings that the volume of statistically distinguishable models is maximal near the critical points of a system and that models far from the critical point are difficult to tell apart using their data^44^.

To summarize these results, when weights are very weak, the occupied state spaces of competing models are essentially as large as the full state space of uncoupled neurons, and thus are fully overlapping and indistinguishable, (Fig. 5f, left). At intermediate weights, the occupied state space shrinks for each model, as does their overlap (Fig. 5f, center). At high weights, the state spaces of the competing models continue to shrink, but instead of growing more distinguishable they shrink toward a common point and their overlap grows again (Fig. 5f, right).

Qualitatively, the distributional convergence of activity states for the competing models shows that it becomes intrinsically difficult, for any algorithm, to distinguish between them at high recurrent weights, consistent with the findings above for various specific inference algorithms. Quantitatively, however, each of the specific inference algorithms suffers from an additional effect when there is model mismatch: because the bias errors do not shrink with data volume but the variance errors do, the best inference, depending on data volume, usually occurs at weaker weights than the point of optimal inherent distinguishibility (were the inference and generative models matched).

### Strong weights greatly exacerbate inference error in partially observed networks

In partially observed networks, it is impossible in principle, without additional assumptions, to infer the existence of a connection between a pair of neurons from observed activity because of the possibility that observed correlations are due to an unobserved common input (e.g. Fig. 1a, if the neuron at the bottom were not observed). To decouple these known problems of inference in partially observed networks ^55–61^ from the more subtle but important problems of inference in recurrent circuits with strong weights, we considered above the fully observed setting. Now we examine how errors that already result from inference in partially observed networks are affected by the strength of weights within the circuit.

As before, we apply a GLM to infer circuit connectivity. This time, we use activity data from only a subset of neurons to reconstruct connectivity within that subset. We repeat this process over multiple subsets of the same size, and “merge” the inferred sub-circuits together (for each neuron pair *i*, *j*, we average the inferred weights *Ŵ*_*ij*_ from all sub-circuits that contained that pair) to obtain a complete *N* × *N* circuit that can be visualized. For fixed, weak weights, as the observed fraction shrinks, the side bands become more pronounced (Fig. 6a, left two panels), illustrating that partial observation leads to qualitatively similar patterns of bias error as when weights are strong. Similarly, for a given observed fraction, increasing the weight strength produces stronger side bands (Fig. 6a, second and fourth panels).

Quantitatively, at fixed data volume the inference error rises linearly as the observed fraction shrinks. Increasing the weight strengths causes an increase in the rate at which circuit inference deteriorates with the observed fraction (Figs. 6b, c, d). In other words, the errors introduced by partial observation are milder when the weights are weak, and bigger when they are strong, even when the architecture is fixed.

The increase in the slope of inference error versus observed fraction when weights are larger is due to mounting bias errors, Fig. 6e. At the weakest weights, inference quality in the 50% observed network nearly matches the fully observed case (Fig. 6f, left), but the gap between the two is large at strong weights (Fig. 6f, right), and the gap, due to bias errors, is largely immune to data volume.

**Figure 6:**
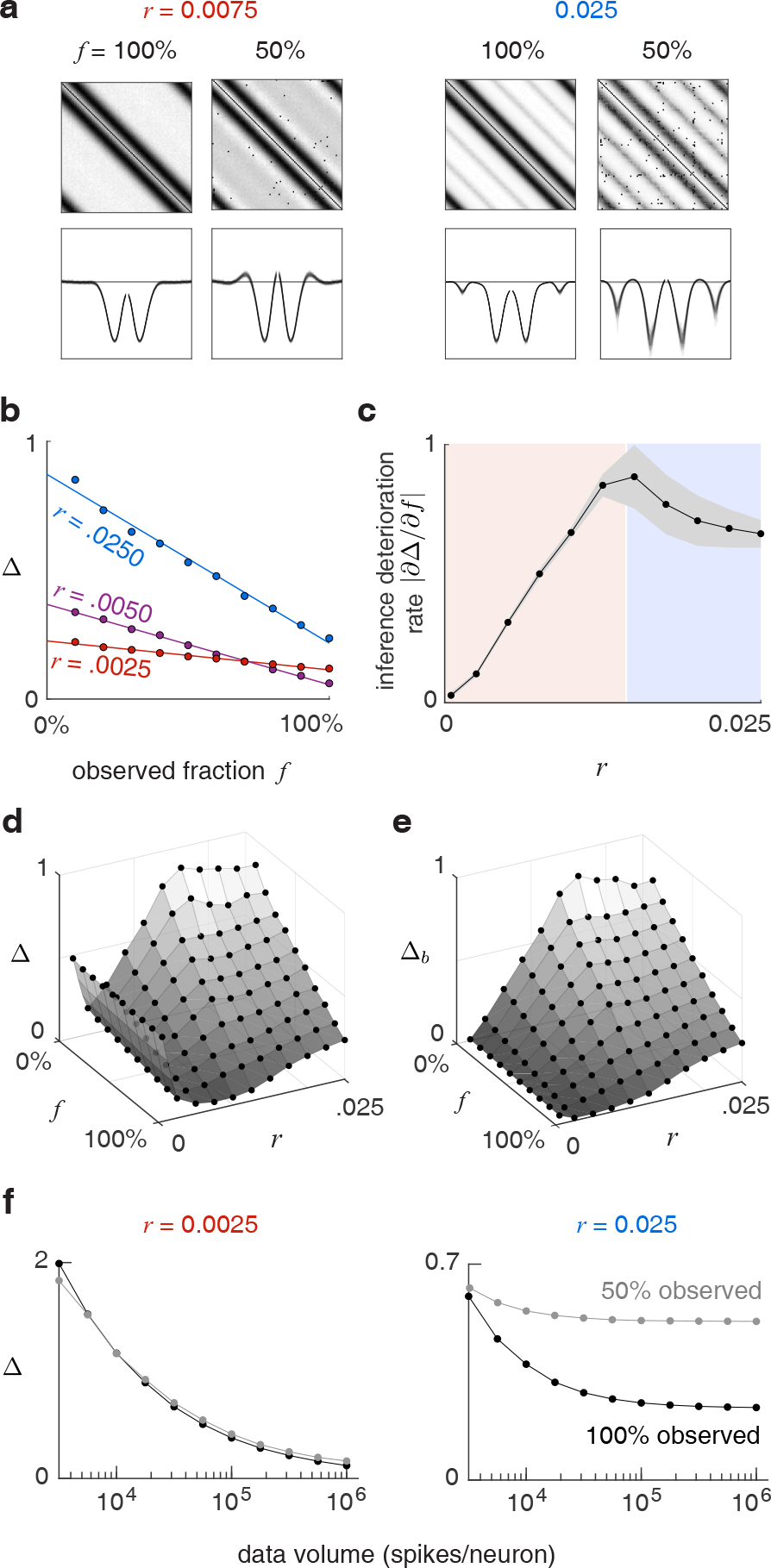
Bias errors in inferring a partially observed circuit are worsened when weights are strong. **a:** Weight matrices (top) and superposition of weight profiles from each node to the rest (bottom), of *merged* inferred sub-circuits, for fully and 50% observed networks, at weak and strong weights. **b:** Average error of inferring sub-circuits of the network of different fractional sizes, at different weight strengths. **c:** The rate of inference quality deterioration with reducing observed fraction (fitted slopes to plot c, with 95% confidence bounds), against weight strength. **d,e:** Full surfaces of average inference error, and average bias error in the merged inferred circuit, against observed fraction and weight strength. Black dots are data points, surfaces are interpolations. **f:** Inference error against data volume for fully and 50% observed networks, at weak and strong weights.

## Discussion

We have shown that estimates of functional or effective connectivity in a network obtained from activity at the nodes can diverge substantially and in systemically biased ways from actual connectivity in a recurrent circuit if the weights are strong. Our definition of strong weights encompasses not only circuits that can hold states without external inputs, as required for memory, but also sensory circuits which moderately or strongly amplify their inputs. Note too that although the issues we illustrate were in the context of inferring connectivity within a network from spike data, they may arise more generally in other domains as well, such as when considering other kinds of activity data or connectivity across networks, such as when inferring area-wise connectivity from fMRI activity data.

The divergence between estimated and true connectivity cannot be addressed by collecting more data because the explaining away errors found in strongly coupled networks are biases that do not shrink with data volume, as do noise-based variance errors which dominate at weak connectivity. Moreover, estimates of functional connectivity by sophisticated inference algorithms in the strongly coupled regime are generally not much better than simple inverse correlation, except in the strongly connected regime where they both provide relatively poor estimates of the true connectivity. These results point to the need for considerable caution when using activity to construct an estimate of structural connectivity in even moderately strongly coupled recurrent circuits.

Arguably, it is sufficient for many purposes to merely infer functional connectivity, without relating it to structural connectivity. This can be the case when the goal is to use activity-based inference to ultimately predict future activity, as done for instance in ^7;8^. However, the functional connectivity in even these cases is then often referred to and used as a proxy for actual connectivity, and here systemic biases in estimation can lead to important errors in our understanding of circuit and circuit development mechanisms. For example, in our simple ring network, purely local connectivity can give rise to periodic activity patterns. One could imagine that developmentally, such a circuit with periodic activity patterns could arise without the need for activity-dependent plasticity. On the other hand, in a sideband model, the sidebands are related to the periodic activity patterns, suggesting that activity must have lead to connectivity, a developmentally distinct process.

Despite the pessimistic results shown here, the challenge of discovering the connectivity in recurrent neural networks is difficult but not unsurmountable: rapid advances in automated segmentation for connectomics mean that one can imagine obtaining connectivity matrices for complete circuits in the not-too-remote future. However, the costs remain large, and determining the link between structure and function after obtaining a connectivity matrix still requires an inference step or model of how activity emerges from connectivity. More immediately accessible and directly interpretable are experiments that rely on perturbation of the system. Clearly, if every single connection could be individually perturbed, that would provide detailed causal connectivity information.

But even much lower-dimensional perturbations can be helpful: Appropriately designed perturbations can help disambiguate between competing dynamical models with even subtle differences^62–64^. More generally, given that variance errors can be averaged away, and that inference with strong noise is superior to inference with weaker noise given enough data (better inference when the noise to recurrent weight strength ratio is high) as we have shown, we expect that driving systems strongly with noise and then performing even simple correlational inference (with lots of data) should lead to much better estimates of true connectivity than when the system is not noise-driven even if sophisticated inference algorithms are used. In other words, studying a recurrent dynamical system with fixed points as it is continuously driven out-of-equilibrium by noise should be much more informative about its structure than studying it close to or at equilibrium ^54^. We have additionally shown above that studying far-from-equilibrium dynamics in strongly coupled recurrent networks, e.g. by watching the network evolve from a fully silenced to patterned state, can provide data for much better inference in strongly recurrent networks.

Finally, when the inference model exactly matches the dynamical model generating the data, all neurons are observed, and the mapping from circuits to activity is injective, it can be possible to exactly estimate connectivity from activity (e.g. in Ising-on-Ising inference for certain architectures). While it is impossible for any inference model to exactly match the true neural circuit dynamics, and the mapping from circuits to activity states need not be injective, improvements in how well the inference models mimic the dynamics of the system can shrink the gap between functional and structure connectivity.

## Acknowledgements

We thank Matthias Bethge, Peter Dayan, Ingmar Kanitscheider, Berk Gerçek and Rishidev Chaudhuri for helpful discussions. Funding was provided by the HHMI through the Faculty Scholars program and the Simons Collaboration on the Global Brain through the Simons Foundation.

## Methods

### Generative network model

Here we outline the dynamical model that was used to generate the data in most cases. Other generative models that we used are described in the Supp. Info.

#### Circuit

Neurons are arranged on a ring. The outgoing synaptic weights *W*_*ij*_ from each neuron to all of the others around the ring have a local Mexican-hat (difference-of-Gaussians) shape:

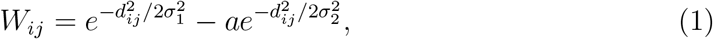

where *d*_*ij*_ is the distance (in neurons) between neurons *i* and *j*. We set *a* = 1.0005 > 1 to make the weights purely inhibitory, to prevent self-excitation and allow dynamical stability. Parameters used: *σ*_1_ = 6.98 and *σ*_2_ = 7 (in neurons).

#### Neural dynamics

Dynamics are updated in discrete time, with a time-step Δ*t* of size 0.1ms. The vector of inputs to the 100 neurons is:

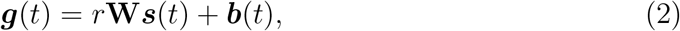

where the synaptic activations of neurons are given by ***s***(*t*), **W** is the recurrent connectivity defined above, and **b** are the feedforward inputs. The relative influence of **W** is scaled by the weight strength *r*. ***b***(*t*) is given by: ***b***(*t*) = *b*(**1** + ***ξ***(*t*)), where *b* = 0.001 provides a uniform excitatory drive, and ***ξ***(*t*) is a multiplicative private Gaussian white noise per neuron, with zero mean and s.d. *σ*_*ξ*_ = 0.3, resulting in a Poisson-like variance proportional to the mean activation. This noise is only injected with probability ≈ .07 in each time step of the discrete-time equations, since with more noise the dynamics loses coherence.

If at any time step (*t*, *t* + Δ*t*] the input *g*_*i*_ to neuron *i* exceeds threshold Θ, it emits a spike. ***σ***(*t*) is the binary vector of spikes from the network at time step (*t*, *t* + Δ*t*]. The synaptic activation of neuron *i*, through which it affects the other neurons in the circuit, is incremented by an amount Δ*t* whenever that neuron spikes, and otherwise decays exponentially according to:

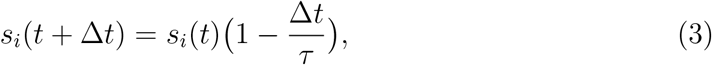

with a synaptic time constant *τ* = 10 ms.

#### Generating spike data

To move from the weakly to the strongly coupled regime, we increase the weight strength *r*. The threshold Θ is adjusted at each *r* to hold roughly fixed (within 16.0 ± 0.1 ms) the average inter-spike interval of the combined across-population spike train.

For each parameter setting, we initialize the network dynamics with random activations ***s*** and wait for it to equlibriate, then collect a total volume of 10^8^ spikes from the network.

### Characterizing the dynamical regime of the network

#### Noise correlations

To measure noise correlations across the network, we need to hold the signal fixed. Removing the feedforward input *b* from a neuron forces a valley of activity at its location, and holds the activity pattern stationary. We then generate spikes from the network with no further time-binning than the generative resolution (which would artificially amplify correlations by calculating them over time). We then calculate the correlation coefficient between the (unbinarized) spike count trains of every neuron pair, excluding the suppressed neuron. Strong positive and negative values both indicate correlated activity, so we take only the magnitude.

#### Activity pattern strength

The phase of the spatially periodic activity pattern randomly wanders around the network over time. But its degree of spatial periodicity will be captured by activity correlations regardless of its phase. We start with spike data that is not binned in addition to the generative resolution of 0.1 ms, and compute the correlation coefficient matrix between all neuron pairs (replacing the 1’s in the diagonal with 0’s to ignore self-correlation). Since the pattern can migrate around the network while the data is being collected, these correspond to signal correlations. A global spatial periodicity of the pattern is reflected as a periodicity in each row/column of this matrix. We calculate the degree of this periodicity in each row by averaging its autocorrelation at circular shifts of 1, 2 and 3 periods (the pattern has 4 periods around the ring), then averaging this across all rows. This measure will be near 0 for uncorrelated spike trains, and approaches 1 in the limit of a 4-period spatial pattern in spike-time correlations.

#### Diffusivity of activity pattern phase

Over time, the phase of the activity pattern migrates randomly around the network. To measure the diffusion coefficient of this motion, we choose the Fourier component of the vector of neural inputs ***g***(*t*) with the spatial frequency of the activity pattern (4 cycles around the ring). The time-varying phase *ϕ*(*t*) of this component is the trajectory of the pattern phase. The slope of the linear fit of 〈(*ϕ*(*t* + *τ*) − *ϕ*(*t*))^2^〉 _*t*_ against *τ* then gives the diffusion coefficient. For weights below *r* = 0.0125, ***g*** was too noisy to extract any periodic pattern, hence diffusivity was not computed for these.

### Measuring inference error

The ground-truth and inferred weights are the elements of **W** and **Ŵ**, ignoring the diagonals. Since the inference model is generally different from the generative model, **Ŵ** has an arbitrary overall scale factor with respect to **W**. So before calculating the inference error, the first step is to re-scale **Ŵ** to match the scale of **W**, which we do in the following way.

**W** is circulant, so each row is a rotation of the same Mexican-hat shaped weight vector ***ω***. Because of noise, though, the rows of **Ŵ** are not exact rotated copies of each other. Each row *i* is a different noisy estimate ***ω***_*i*_ of ***ω***. We re-scale **Wˆ** such that its average weight-shape estimate 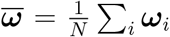 has the least *l*_1_ deviation from the true shape ***ω***. Averaging the noisy ***ω***_*i*_’s and using the *l*_1_ metric (which is more noise-tolerant than *l*_2_) provide greater accuracy in the scale-matching in the face of noise.

However, this procedure cannot be applied in situations where the weight matrix to be inferred is not circulant. This is the case when inferring a partially observed circuit, (a sub-matrix of **W**, see Results, Fig. 6a,b), and also when inferring a circuit with a non-circulant, e.g. random architecture (see Results, Fig. 4c). In these cases we resort to the less noise-tolerant choice of re-scaling the inferred matrix **Ŵ** to have the least *l*_2_ distance from the ground-truth matrix **W**. When we apply this re-scaling, the inference error is also the sine of the angle between the ground-truth and inferred weight vectors in the space of weight parameters.

After **Ŵ** has been re-scaled, the inference error is the magnitude of the (*l*_2_) distance between the vectors of ground-truth and inferred weights, expressed as a fraction of the vector magnitude of the ground-truth weights:

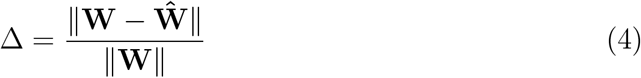

Although the *l*_2_ metric is more noise-sensitive as noted, we choose it here in the final expression, since its properties allow an elegant decomposition of the inference error into variance and bias components, as we describe in the next section.

#### Variance and bias errors

Let us denote the *N*-1 elements (ignoring self-coupling) of the common weight vector ***ω*** of the circulant **W** by *ω*^*α*^, and their estimates on row *i* of **Ŵ** by 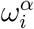. The squared inference error (eq. 4) can thus be expressed as:

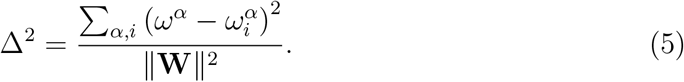

This can be rewritten in terms of 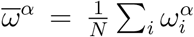, the elements of the average estimated weight shape 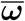:

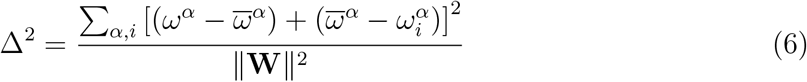

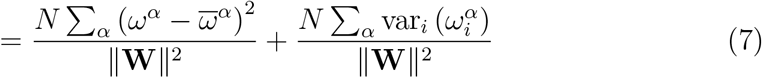

(where 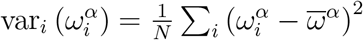 is the variance of the estimates of *ω*^*α*^)

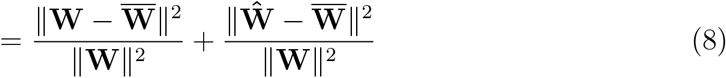

(where 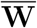 is a circulant matrix consisting of rotations of the mean estimate 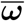)

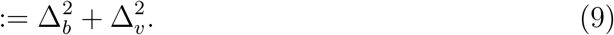

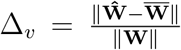, the *variance error*, is the error due to the variance among the different noisy estimates 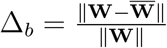, the *bias error*, is the error due to deviation of the mean estimate 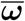 from the true weights *ω*. We can geometrically interpret these quantities in the space of all *N* (*N* − 1) weight parameters. **W**, **Ŵ** and 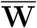 are three points in this space representing the true weights, inferred weights, and mean inferred weights. If we measure lengths in units of ‖**W**‖, the length of the gap between **Ŵ** and **W** is the total inference error Δ, and can be decomposed into the gap due to variance, from **Ŵ** to 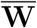 with length Δ_*v*_, and the gap due to bias, from 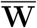 to **W** with length Δ_*b*_. These two gaps are orthogonal, as evidenced by their Pythagorean relationship (eq. 9), and their zero inner product:

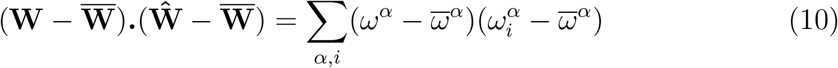

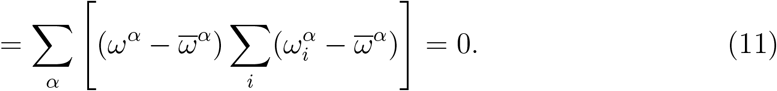

Hence, the total inference error is a vector sum of orthogonal components due to variance and bias.

The relative contribution of bias to the inference error can thus be measured by the angle between the vectors of the total inference error and the variance error, as a fraction of 90°:

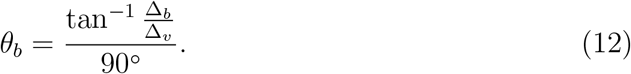

These variance and bias errors cannot be computed for non-circulant circuits.

### Discriminating circuits using activity data

#### Constructing the alternative circuit

The circuit inferred at strong weights using inverse Ising needs to modified to be used as a generative circuit with the same dynamics as our original generative model.

We first calculate the mean weight shape vector 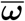 of this inferred circuit by circularly shifting the matrix rows to align them, then averaging. Next, we set the positive parts of this mean shape to 0 (to prevent self-excitation and ensure stability under the generative dynamics), and re-scale to match its minimum with that of the ground-truth weight shape. Finally, we create a circulant matrix from this weight shape.

As with the original circuit, firing thresholds Θ for this circuit are tuned at each weight strength to set the average network inter-spike-interval at 16.0 ± 0.1 ms. We can now generate spike data from this circuit using the original dynamics.

#### Selecting a well-sampled subspace of neural activity

Even with binary spike data, there are 2^*N*^ possible spike words, or states, of an *N*-neuron network, prohibitively big for large *N* (in our case with *N* = 100, it is 2^100^ 10^30^). Each time-bin of spikes we collect from the network is a state. The number of such states we obtain over 6 hours at a resolution of 0.1 ms, and binned at 10 ms, is only of the order of 10^6^. Thus, even if in a given setting the network may produce only a subset of all possible states, they are still severely undersampled by data collected in any experiment of reasonable duration, and statistics to characterize the data distribution, such as entropy, will be biased^65;74^.

We can instead choose a segment of *n* adjacent neurons from our ring network, whose smaller *n*-dimensional state distribution we can make sure to have sampled well with our data. But this fails to utilize the data we collected from the other neurons. We can address this by exploiting the rotation-invariance of our network, and its activity over time. Sliding our *n*-neuron window around the ring network, we can pool together the observed spike vectors of neurons 1 through *n*, 2 through *n* + 1 etc. This lets us use the full dataset to build a better-sampled distribution of adjacent *n*-neuron states of the network.

*n* should be chosen as large as possible while ensuring that its state distribution is sampled sufficiently in our data. This largest possible value is dictated by the particular statistical measure we want to accurately estimate, as explained in the following sections.

#### Characterizing neural activity using information theory

##### Entropy

We can verify that a distribution has been well-sampled for unbiased entropy estimation by checking that the entropies computed with increasing fractions of the data converge as we approach the total data volume. *n* = 22 is the largest value that allows this convergence at all weights (see Supp. Fig. S7). However, as described in the following sections, requirements for other measures constrained *n* to be lower.

##### Relative entropy

The dissimilarity between the data distributions *p***W** and *p***W**′, considering the former as the ‘true’ distribution, is measured by their relative entropy (KL divergence):

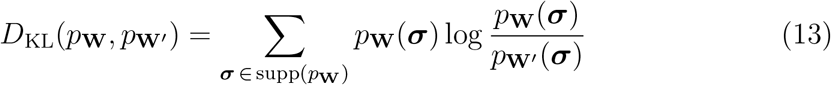

where ***σ*** runs over the *n*-neuron spike vectors that constitute the support of *p***W**.

##### Fisher information

This can be used to quantify the amount of information about a generative model that is contained in the data it produces. It is measured by the sensitivity of the data distribution to changes in the model parameters.

It is not feasible to compute the Fisher information against variations of all the circuit parameters (the 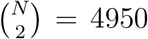 unique weights in **W**). Therefore, we consider instead a single-parameter family of models that passes through the true circuit **W** and the non-local circuit **W**′:

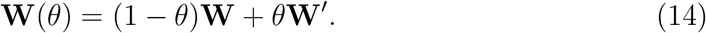

The Fisher information *I*(*θ* = 0) about the true model **W** = **W** (*θ* = 0) can be written in terms of the KL divergence between the data distribution *p*_**W**_ (= *p*_0_) that it produces and the distribution *p*_*dθ*_ that a neighbouring model **W** (*dθ*) produces:

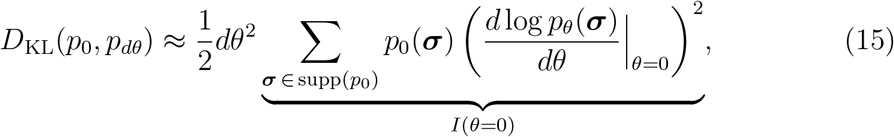

where ***σ*** again constitute *n*-length binary spike vectors.

We approximate this measure by constructing the circuit 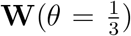 as a neigh-bouring model in the family. At each weight strength, we again adjust its firing threshold to maintain an average network inter-spike interval of 16.0 ± 0.1 ms as before, generate spike data, binarize counts, and compute the KL divergence with respect to the data distribution generated by **W**. The Fisher information is then approximated by:

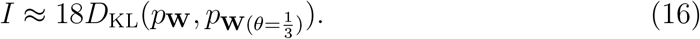

##### Likelihood ratio of circuit models

The log likelihood that the observed data distribution *p*_**W**_ was produced by the circuit **W**′ is:

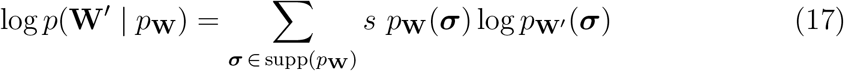

where *s* is the total number of samples in *p*_**W**_.

We can collect a second sample 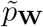 from the local circuit to account for sample-to-sample variability, and then calculate the log likelihood ratio of **W** vs **W**′, given data generated from **W**:

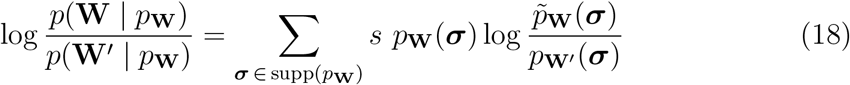

In order for the relative entropy, Fisher information and likelihood ratio to be defined, the distributions must be sampled well enough that each spike state ***σ*** that occurs in *p*_**W**_ occurs in 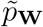, *p*_**W**′_ and 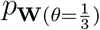. With the total volume of spike data that we collected from the network, the largest value of *n* for which this was possible is 10. Thus, these distributions are over 2^10^ 10-neuron binary spike state vectors.

### Inferring a partially observed network

For each of a range of sub-population sizes *n*, we randomly select multiple *n*-neuron sub-populations from the 100-neuron network. We use the spike data from each such sub-population to infer its corresponding *n×n* connectivity submatrix.

In order to have statistically informative results, for a given sub-population size *n*, we want to choose enough sub-networks such that together they cover the whole network reasonably well. Each *n*-neuron sub-network covers a fraction 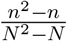 of the weight matrix of the entire network (ignoring diagonal terms). Using this we calculate that randomly selecting

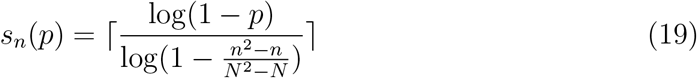

sub-networks ensures with probability *p* that all synapses of the full network have been sampled at least once. We collect enough samples to have *p* = 0.99 for each sub-population size *n*.

## Supplementary information

### S.1 Inference with matched models

All our results are obtained for the realistic case where the (statistical) inference models differs from the (dynamical) model in which the spike data are actually collected. This case of model mismatch is inevitable when one is inferring neural circuitry from experimental recordings, since the full biological dynamics that generate the observed activity are unknown, and arguably more complex than any theoretical model can describe.

Nevertheless, it is useful to complete our understanding of circuit inference by comparing the idealized theoretical case in which the inference model exactly matches the generative model.

We would like to perform inference on our generative dynamical model (see Methods) by using it also as the inference model, except that it is not simple or end-to-end differentiable, so cannot readily be used for inference. Instead, we use the same circuitry **W** that we have used so far, but the simpler Ising model for both the generation of (binary) spike data (Supp. Info. §S.2), and inference (with the minimum probability flow algorithm).

Inference with this matched model is better in several respects (Fig. S1). Comparing the inference error curve with the pattern strength curve (the Ising model has no notion of time, so diffusivity cannot be computed) shows that optimal inference is now just inside the memory regime. Unlike the previous cases, inference errors are variance errors, which decay with data volume as Δ^2^ ~ 1*/D* at all weights, as seen in the uniform drop in the log-log axes of Fig. S1a, secondary axis (see Supp. Fig. S4 for power-law fits). Consequently, the optimal inference point remains stationary with increasing data, reflecting an intrinsic critical point of the system.

These findings are qualitatively reproduced with a model-matched inference using a generalized linear model for both generation and inference (Supp. Info. §S.7). Like Ising-on-Ising inference, inference error is almost entirely due to variance at all weights. Optimal inference is again just inside the memory regime at the point of pattern onset, and remains stationary there as increasing data volume erodes variance error everywhere.

**Figure S1:**
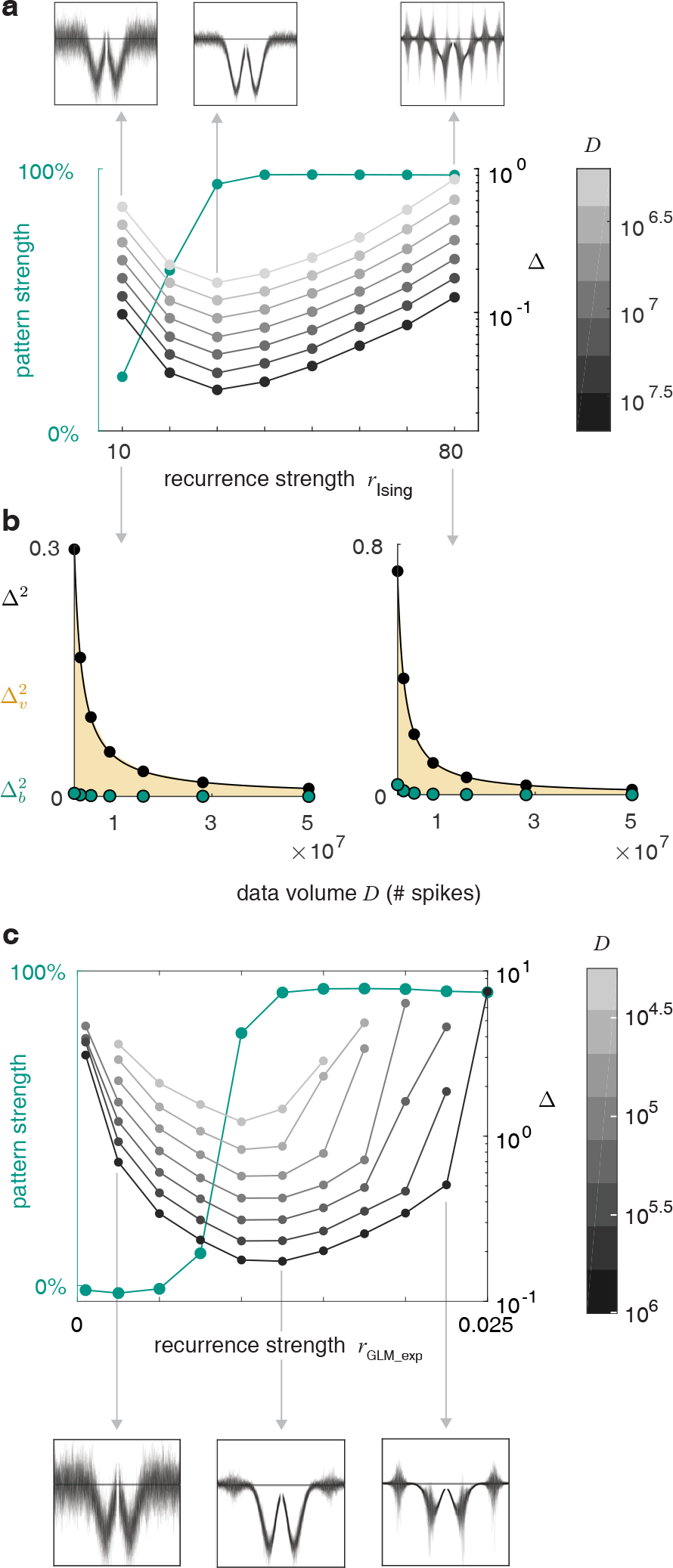
Inference with matched models. **a:** Using an Ising model for both generation and inference. Top: superposed inferred weights from each node to the rest (line marks zero). Bottom: pattern strength, and inference error with different data volumes, against weight strength. **b:** Squared total, variance and bias errors against data volume at weak and strong weights. **c:** Using a generalized linear model with an exponential nonlinearity (§S.7) for both generation and inference. Top: Pattern strength, and inference error with different data volumes, against weight strength. Bottom: Superposed inferred weights from each node to the rest.

### S.2 The Ising model for data generation

To use an Ising model as a generative model, we set the Ising coupling matrix to **J** = *r*_Ising_**W**, where **W** is the same ground-truth weight matrix as used so far, and *r*_Ising_ was varied from 10 to 80 to be in the right physical regime containing the inference optimum. The biases of the Ising model are taken to be uniform: *h*_*i*_ = 1. The Ising states were generated using a Gibbs sampling algorithm that updates the spins in a random sequence in each pass.

### S.3 The Ising model for inference

The Ising model, when used for circuit inference, performs better when the spike data are binned at an appropriate time-scale. The Ising model also requires spike data to be binary. So to use inverse Ising inference, we bin the spike data with different bin-widths and binarize the bins (spike/no spike) to obtain a total of 5 million binned and binarized spikes, then implement inverse Ising inference with minimum probability flow and compute the inference error. We then choose the bin-width that yields the minimum inference error (Supp. Fig. S2b). We use the same procedure to find the best bin-width for inference using logistic regression. The optimal bin-width turns out to approximately equal *τ*, the neural time constant in the generative model. This is consistent since *τ* is the time scale over which the spiking of one neuron can influence that of a connected neuron, hence choosing it as the bin-width bins causally related spikes together, leading to more accurate correlations (Supp. Fig. S2a) and better inference.

Binarizing the spike data for inverse Ising inference results in discarding spikes in bins containing multiple spikes. With stronger weights, the peaks of the activity pattern become sharper and higher; at the peaks of the activity pattern, more bins contain multiple spikes, and more spikes must be discarded (Supp. Fig. S2c). Thus, to implement inverse Ising inference on a total of 10^8^ spikes across weights (as with other inference methods, see Methods), the actual number of spikes collected from the network prior to binarization was larger.

### S.4 Mean-field Ising models

Mean-field Ising models are a fast way to infer weights from binarized spikes, but their simplifying assumptions produce different degrees of approximate solutions depending on the model being studied ^67^. The following are expressions for the weights with the naïve mean field (NMF), which we also call ‘raw correlations’ (*C* is the spike-train covariance matrix and *m*_*i*_ is the average state of the *i*^th^ node):

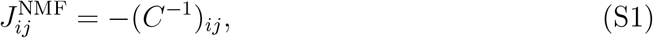

the Thouless-Anderson-Palmer (TAP) model:

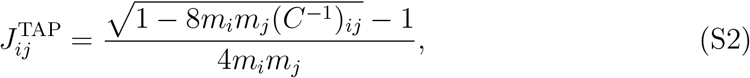

and the Sessak and Monasson (SM) model:

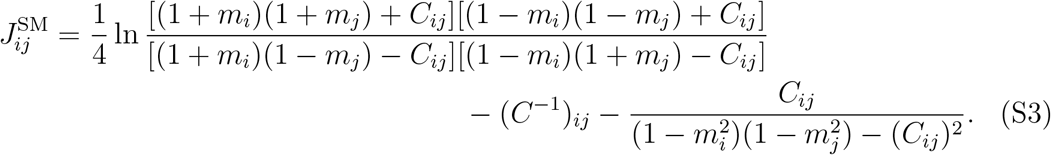

### S.5 Linear-nonlinear-Poisson model

A linear-nonlinear-Poisson (LNP) network was used to generate data from the circuit **W**, for inference with a generalized linear model (Supp. Fig. S6). Its neural inputs are the same as the original dynamics model (see Methods):

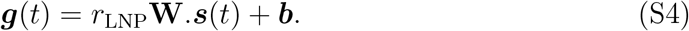

However, here the external input ***b*** is 0.001 everywhere and lacks any explicit noise component. Instead, the input is passed through a rectifying filter to obtain a firing rate. The number of spikes *n*_*i*_(*t*) of each neuron *i* in each time-bin are then Poisson-distributed with this rate:

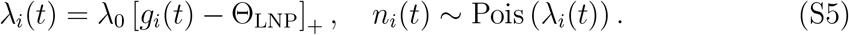

The outgoing activations ***s*** follow the same dynamics as before:

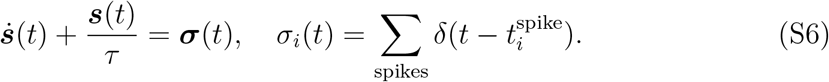

In order to maintain an average network inter-spike interval of 16.0 ± 0.1 ms at each weight strength *r*_LNP_, *λ*_0_ was set to 32, and the threshold Θ_LNP_ was adjusted for each weight strength. The spike data were not further binned for use in GLM-based inference.

### S.6 Generalized linear model (rectified)

This generative model is a close cousin of the LNP model, which is consistent with the fact that inference on them yield similar weight profiles (see Supp. Fig. S6).

The neural inputs are given by a spatiotemporal filter over spikes:

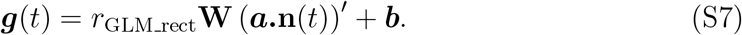

Here **n** (*t*) is an *T* × *N* matrix of the spike-count vectors of the network in the previous *T* = 200 time-bins. ***a*** is a length-*T* exponential kernel that filters the spike history **n** (*t*):

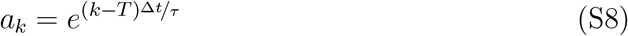

where Δ*t* = 0.1 ms is the resolution at which the discrete-time equations are evolved, and *τ* = 10 ms. This temporal filtering serves a similar puprose to the dynamics of ***s*** in the LNP model. ***b*** is once again 0.001 at every node. As in the LNP model, these inputs are passed through a rectifying inverse link function to produce Poisson firing rates:

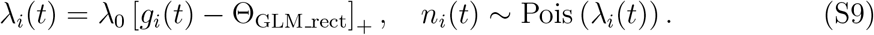

Once again, in order to maintain an average network ISI of 16.0 ± 0.1 ms, *λ*_0_ was set to 100 here, and the threshold Θ_GLM_rect_ was adjusted for each weight strength.

### S.7 Generalized linear model (exponentiated)

A generalized linear model with an exponential nonlinearity (i.e., the canonical logarithmic link function for the Poisson distribution) was used both as a generative and an inference model.

The inputs in this model are the same spatiotemporal filters over the spikes as in the previous model. However, they are then passed through an exponential inverse link function to produce Poisson firing rates:

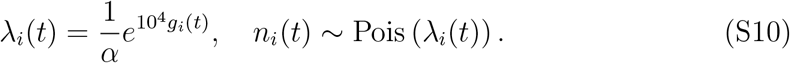

*α* was adjusted at each weight strength to maintain the average network ISI at 16.0 ± 0.1 ms.

### S.8 *l*_1_-regularized logistic regression

When performing logistic regression, we first mean-subtract all the spike channels. Then we binarize the spike counts of the dependent channel, but leave the predictor channels unbinarized. The matrix of regression coefficients of each node against all others is taken to be the inferred weight matrix. At each weight strength we find the regularization strength *λ* that minimizes the inference error (Supp. Fig. S5).

### S.9 Negentropy of inference errors

Once we scale-match **Ŵ** with **W** (see Methods), the elements of **W** − **Ŵ** give us the distribution of errors in individual inferred weights. Negentropy is a way to characterize this error distribution in terms of its dissimilarity from a normal distribution (which would result from purely random errors). If the error distribution has variance *σ*^2^, and *p*_*i*_ denotes the relative frequencies in the distribution histogram binned with binsize *b*, negentropy is given by:

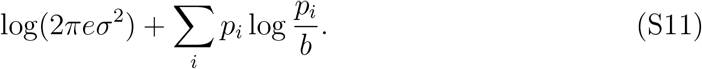

This is the difference between the differential entropy of a normal distribution with the same variance, and a continuous version of the discrete entropy of the error distribution.

**Figure S2:**
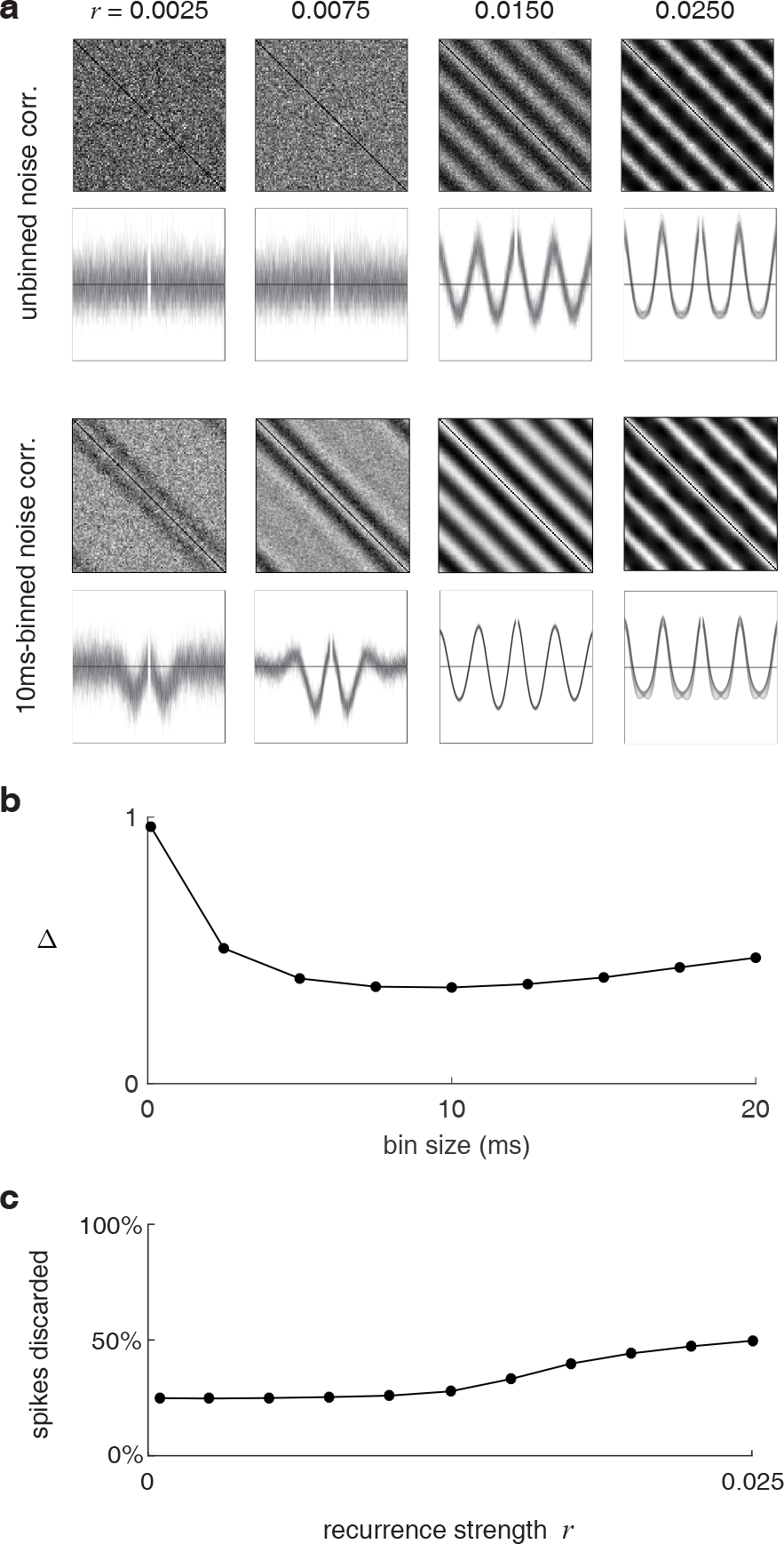
Binning affects noise correlations and inference. **a:** Noise correlations between neuron pairs (top: full matrix, bottom: superposed vectors between each node and the rest) for binned vs unbinned spikes. The right bin-width groups causally related spikes together, and noise correlations at an intermediate *r* then reflect the underlying weights. **b:** Inference error using inverse Ising with MPF on spike data binned at different widths. **c:** Fraction of spikes discarded when binarizing binned spike data for Inverse Ising inference.

**Figure S3:**
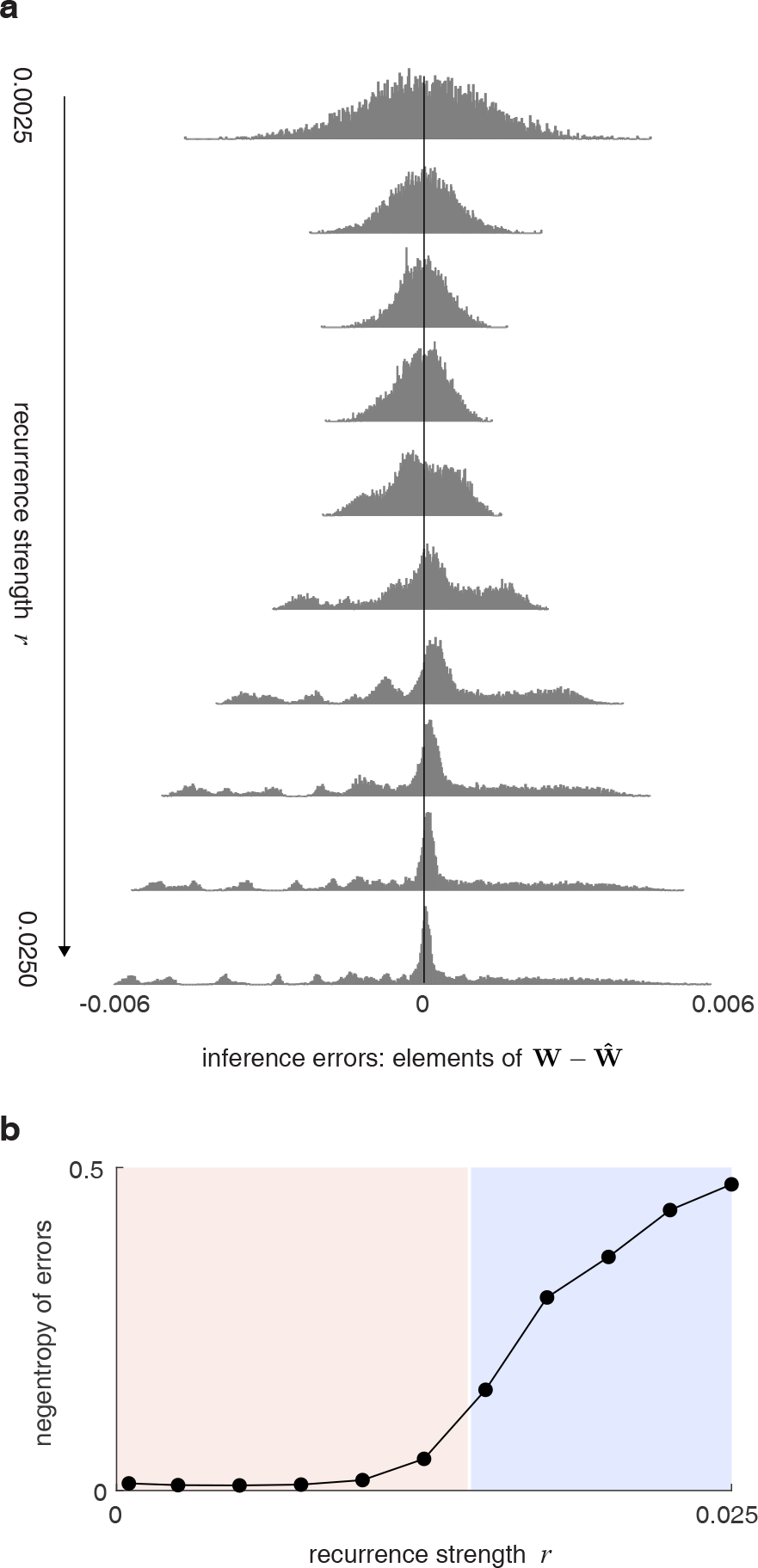
**a:** Distribution of errors in inferred connectivity (relative to the length of the ground-truth weight vector) at different weights. With weak weights, errors are random, thus normally distributed. As the weight increases, errors initially shrink as noise weakens and SNR grows; the distribution of errors becomes increasingly non-normal with increasing weight due to bias. **b:** Negentropy of the error distribution (§S.9) against weight strength.

**Figure S4:**
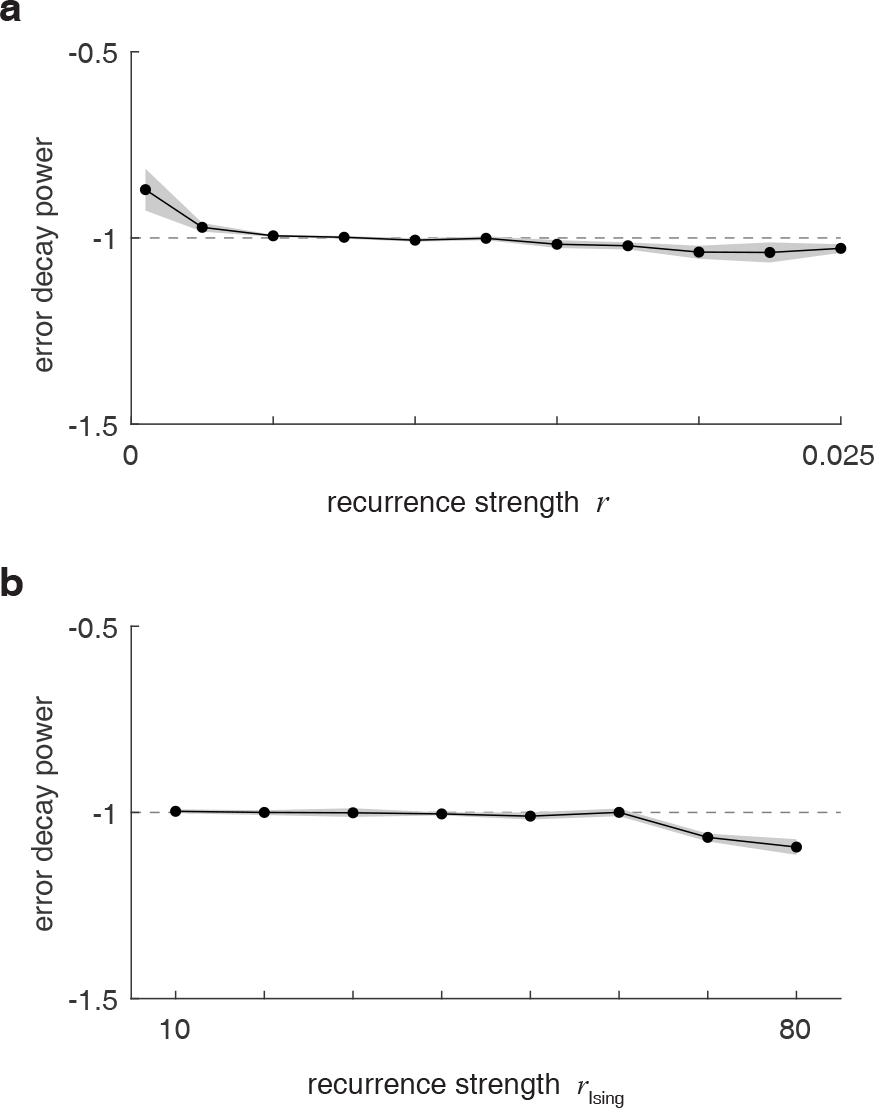
**a:** The fitted exponents *α* of the power-law decay of variance error with data volume 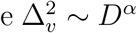 when using a generalized linear model to infer on data from the original model. Error-bands are 95% confidence intervals. The theoretical exponent is −1. **b:** The exponent *α* of the power-law decay of total inference error with data volume Δ^2^ ∼ *D*^*α*^ when using the Ising model for both data generation and inference. Here inference error is almost entirely due to variance, thus follows the power-law decay with data.

**Figure S5:**
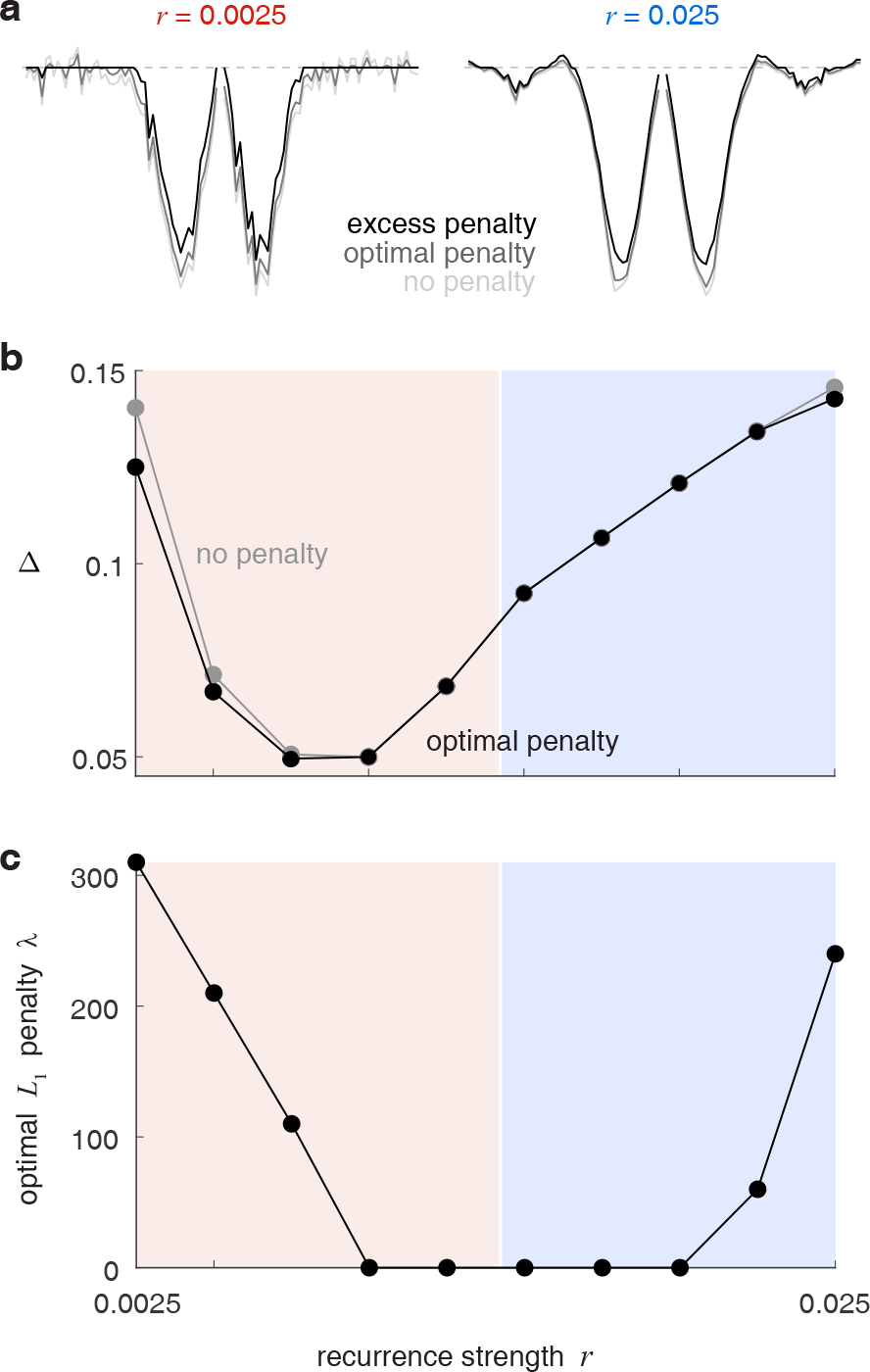
Circuit inference using logistic regression is not improved by *l*_1_ regularization. **a:** Example coupling profiles inferred using logistic regression with zero, optimal and excessive regularization penalties. When weights are weak, regularization reduces some noise and marginally improves inference. At high weights, regularization suppresses both the spurious side-bands and the true coupling shape, so is not helpful. **b:** Inference error vs weight strength using logistic regression with and without *l*_1_ regularization. **c:** Optimal *l*_1_ penalties (that produce the lowest inference errors) at each weight. Regularization barely improves inference in the strong and weak weight regimes.

**Figure S7:**
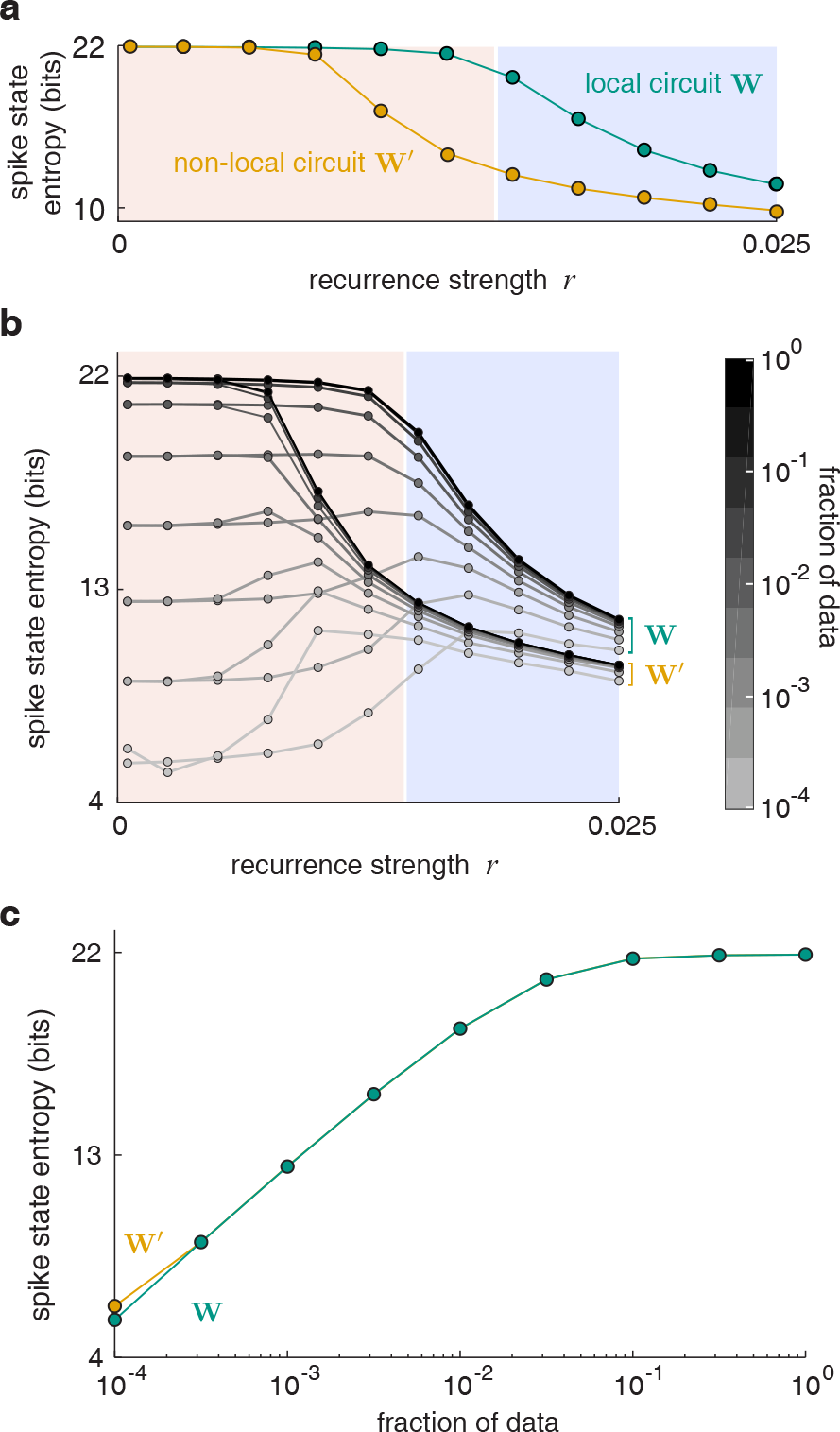
**a:** Entropy of the distributions of 22-neuron spike sub-states from the local circuit **W** and the non-local circuit **W**′. **b:** Entropies of the spike sub-states of the two circuits computed with different data fractions across weight strengths. At all weights, the computed entropies converge as the data approaches the total volume. **c:** Example slice of plot b at the weakest weights, where entropy convergence takes the longest.

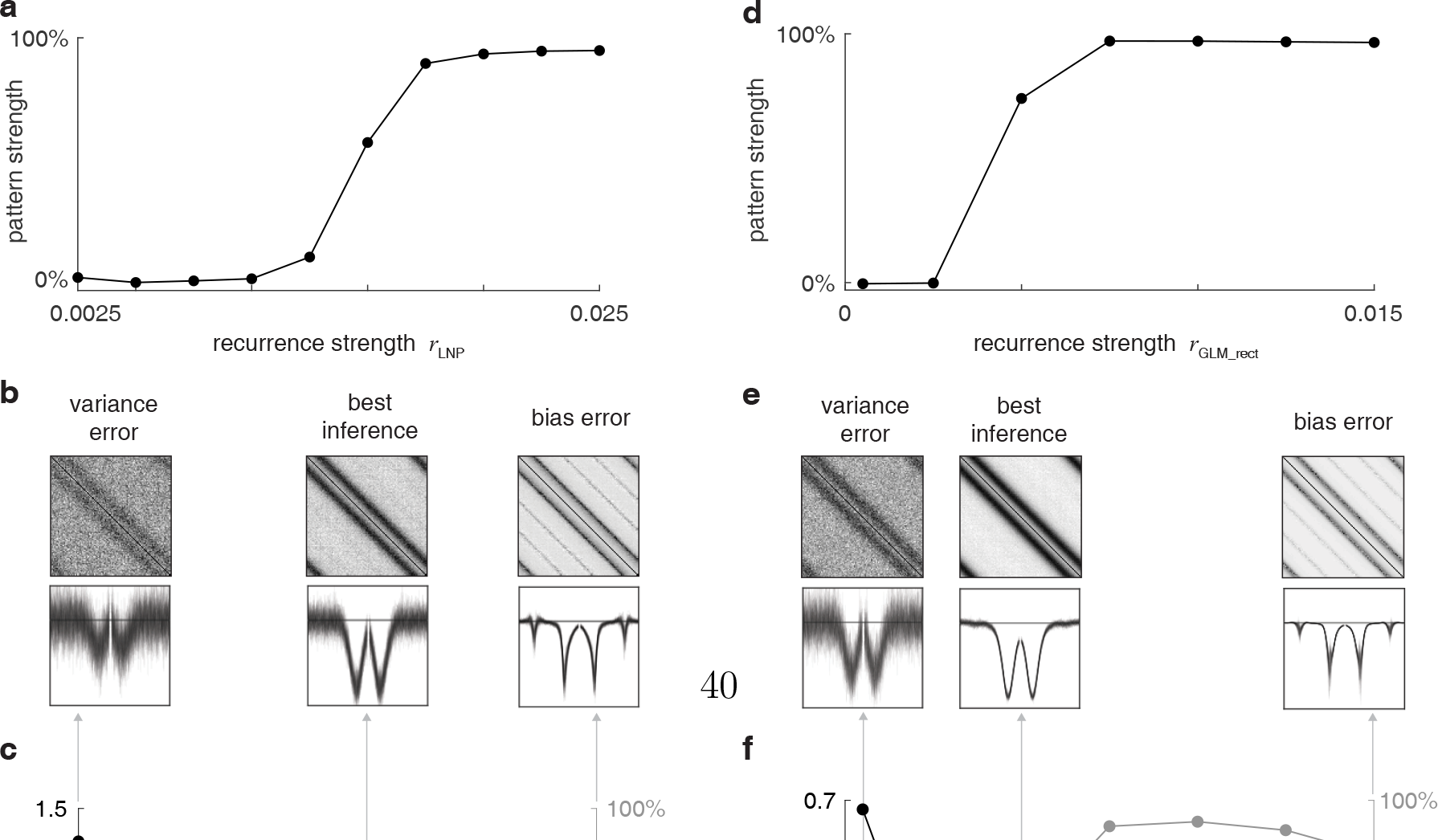

## References

[1] Shin-ya Takemura, Arjun Bharioke, Zhiyuan Lu, Aljoscha Nern, Shiv Vitalade-vuni, Patricia K Rivlin, William T Katz, Donald J Olbris, Stephen M Plaza, Philip Winston, et al. A visual motion detection circuit suggested by drosophila connectomics. Nature, 500(7461):175–181, 2013.

[2] Moritz Helmstaedter, Kevin L Briggman, Srinivas C Turaga, Viren Jain, H Se-bastian Seung, and Winfried Denk. Connectomic reconstruction of the inner plexiform layer in the mouse retina. Nature, 500(7461):168, 2013.

[3] H Sebastian Seung. Continuous attractors and oculomotor control. Neural Net-works, 11(7):1253–1258, 1998.

[4] David Kleinfeld, Arjun Bharioke, Pablo Blinder, Davi D Bock, Kevin L Brig-gman, Dmitri B Chklovskii, Winfried Denk, Moritz Helmstaedter, John P Kaufhold, Wei-Chung Allen Lee, et al. Large-scale automated histology in the pursuit of connectomes. Journal of Neuroscience, 31(45):16125–16138, 2011.

[5] Randal Burns, Kunal Lillaney, Daniel R Berger, Logan Grosenick, Karl Deis-seroth, R Clay Reid, William Gray Roncal, Priya Manavalan, Davi D Bock, Narayanan Kasthuri, et al. The open connectome project data cluster: scalable analysis and vision for high-throughput neuroscience. In Proceedings of the 25th International Conference on Scientific and Statistical Database Management, page 27. ACM, 2013.

[6] Wei-Chung Allen Lee, Vincent Bonin, Michael Reed, Brett J Graham, Greg Hood, Katie Glattfelder, and R Clay Reid. Anatomy and function of an excitatory network in the visual cortex. Nature, 532(7599):370, 2016.

[7] Jonathan W Pillow, Jonathon Shlens, Liam Paninski, Alexander Sher, Alan M Litke, EJ Chichilnisky, and Eero P Simoncelli. Spatio-temporal correlations and visual signalling in a complete neuronal population. Nature, 454(7207):995–999, 2008.

[8] Elad Schneidman, Michael J Berry, Ronen Segev, and William Bialek. Weak pairwise correlations imply strongly correlated network states in a neural population. Nature, 440(7087):1007–1012, 2006.

[9] arl J Friston. Functional and effective connectivity: a review. Brain connectivity, 1(1):13–36, 2011.

[10] Ari Pakman, Jonathan Huggins, Carl Smith, and Liam Paninski. Fast state-space methods for inferring dendritic synaptic connectivity. Journal of computational neuroscience, 36(3):415–443, 2014.

[11] Daniel A Dombeck, Christopher D Harvey, Lin Tian, Loren L Looger, and David W Tank. Functional imaging of hippocampal place cells at cellular resolution during virtual navigation. Nature neuroscience, 13(11):1433–1440, 2010.

[12] Yaniv Ziv, Laurie D Burns, Eric D Cocker, Elizabeth O Hamel, Kunal K Ghosh, Lacey J Kitch, Abbas El Gamal, and Mark J Schnitzer. Long-term dynamics of ca1 hippocampal place codes. Nature neuroscience, 16(3):264–266, 2013.

[13] James J Jun, Nicholas A Steinmetz, Joshua H Siegle, Daniel J Denman, Marius Bauza, Brian Barbarits, Albert K Lee, Costas A Anastassiou, Alexandru Andrei, Çağatay Aydın, et al. Fully integrated silicon probes for high-density recording of neural activity. Nature, 551(7679):nature24636, 2017.

[14] John J Hopfield. Neural networks and physical systems with emergent collective computational abilities. Proceedings of the national academy of sciences, 79(8):2554–2558, 1982.

[15] John Hertz, Anders Krogh, and Richard G Palmer. Introduction to the theory of neural computation. Addison-Wesley/Addison Wesley Longman, 1991.

[16] Daniel J Amit. Modeling brain function: The world of attractor neural networks. Cambridge university press, 1992.

[17] Gasper Tkacik, Elad Schneidman, II Berry, J Michael, and William Bialek. Ising models for networks of real neurons. arXiv preprint q-bio/0611072, 2006.

[18] Pradeep Ravikumar, Martin J Wainwright, John D Lafferty, et al. High-dimensional Ising model selection using l1-regularized logistic regression. The Annals of Statistics, 38(3):1287–1319, 2010.

[19] P McCullagh and JA Nelder. Generalised linear modelling. Chapman and Hall, London. Negro, JJ & Hiraldo, F. (1992) Sex ratios in broods of the lesser kestrel Falco naumanni. Ibis, 134:190–191, 1983.

[20] Liam Paninski. Maximum likelihood estimation of cascade point-process neural encoding models. Network: Computation in Neural Systems, 15(4):243–262, 2004.

[21] Wilson Truccolo, Uri T Eden, Matthew R Fellows, John P Donoghue, and Emery N Brown. A point process framework for relating neural spiking activity to spiking history, neural ensemble, and extrinsic covariate effects. Journal of neurophysiology, 93(2):1074–1089, 2005.

[22] Yuriy Mishchencko, Joshua T Vogelstein, and Liam Paninski. A Bayesian approach for inferring neuronal connectivity from calcium fluorescent imaging data. The Annals of Applied Statistics, pages 1229–1261, 2011.

[23] Duane Q Nykamp. Reconstructing stimulus-driven neural networks from spike times. In Advances in Neural Information Processing Systems, pages 325–332, 2003.

[24] Jonathon Shlens, Greg D Field, Jeffrey L Gauthier, Matthew I Grivich, Dumitru Petrusca, Alexander Sher, Alan M Litke, and EJ Chichilnisky. The structure of multi-neuron firing patterns in primate retina. The Journal of neuroscience, 26(32):8254–8266, 2006.

[25] Eizaburo Doi, Jeffrey L Gauthier, Greg D Field, Jonathon Shlens, Alexander Sher, Martin Greschner, Timothy A Machado, Lauren H Jepson, Keith Math-ieson, Deborah E Gunning, et al. Efficient coding of spatial information in the primate retina. Journal of Neuroscience, 32(46):16256–16264, 2012.

[26] Yuriy Mishchencko, Joshua T Vogelstein, and Liam Paninski. A bayesian approach for inferring neuronal connectivity from calcium fluorescent imaging data. The Annals of Applied Statistics, pages 1229–1261, 2011.

[27] Yuriy Mishchenko and Liam Paninski. A bayesian compressed-sensing approach for reconstructing neural connectivity from subsampled anatomical data. Journal of computational neuroscience, 33(2):371–388, 2012.

[28] Ben Shababo, Brooks Paige, Ari Pakman, and Liam Paninski. Bayesian inference and online experimental design for mapping neural microcircuits. In Advances in Neural Information Processing Systems, pages 1304–1312, 2013.

[29] Alexandro D Ramirez and Liam Paninski. Fast inference in generalized linear models via expected log-likelihoods. Journal of computational neuroscience, 36(2):215–234, 2014.

[30] Diego A Gutnisky and Valentin Dragoi. Adaptive coding of visual information in neural populations. Nature, 452(7184):220–224, 2008.

[31] Jasper Poort and Pieter R Roelfsema. Noise correlations have little influence on the coding of selective attention in area V1. Cerebral Cortex, 19(3):543–553, 2009.

[32] Jason M Samonds, Brian R Potetz, and Tai Sing Lee. Cooperative and competitive interactions facilitate stereo computations in macaque primary visual cortex. Journal of Neuroscience, 29(50):15780–15795, 2009.

[33] Marlene R Cohen and John HR Maunsell. Attention improves performance primarily by reducing interneuronal correlations. Nature neuroscience, 12(12):1594–1600, 2009.

[34] Jude F Mitchell, Kristy A Sundberg, and John H Reynolds. Spatial attention decorrelates intrinsic activity fluctuations in macaque area V4. Neuron, 63(6):879–888, 2009.

[35] Xin Huang and Stephen G Lisberger. Noise correlations in cortical area MT and their potential impact on trial-by-trial variation in the direction and speed of smooth-pursuit eye movements. Journal of Neurophysiology, 101(6):3012–3030, 2009.

[36] Marlene R Cohen and William T Newsome. Context-dependent changes in functional circuitry in visual area MT. Neuron, 60(1):162–173, 2008.

[37] Ehud Zohary, Michael N Shadlen, and William T Newsome. Correlated neuronal discharge rate and its implications for psychophysical performance. Nature, 370(6485):140, 1994.

[38] Wyeth Bair, Ehud Zohary, and William T Newsome. Correlated firing in macaque visual area MT: time scales and relationship to behavior. Journal of Neuroscience, 21(5):1676–1697, 2001.

[39] Caleb Kemere, Margaret F Carr, Mattias P Karlsson, and Loren M Frank. Rapid and continuous modulation of hippocampal network state during exploration of new places. PloS one, 8(9):e73114, 2013.

[40] Xaq Pitkow and Markus Meister. Decorrelation and efficient coding by retinal ganglion cells. Nature neuroscience, 15(4):628, 2012.

[41] Ingmar Kanitscheider, Ruben Coen-Cagli, and Alexandre Pouget. Origin of information-limiting noise correlations. Proceedings of the National Academy of Sciences, 112(50):E6973–E6982, 2015.

[42] Rishidev Chaudhuri, Berk Gerek, Biraj Pandey, Adrien Peyrache, and Ila R Fiete. Dynamics in a canonical microcircuit. Submitted, 2018.

[43] Yoram Burak and Ila R Fiete. Fundamental limits on persistent activity in networks of noisy neurons. Proceedings of the National Academy of Sciences, 109(43):17645–17650, 2012.

[44] Iacopo Mastromatteo and Matteo Marsili. On the criticality of inferred models. Journal of Statistical Mechanics: Theory and Experiment, 2011(10):P10012, 2011.

[45] Tamara Broderick, Miroslav Dudik, Gasper Tkacik, Robert E Schapire, and William Bialek. Faster solutions of the inverse pairwise ising problem. arXiv preprint arXiv:0712.2437, 2007.

[46] Francisco Barahona. On the computational complexity of Ising spin glass models. Journal of Physics A: Mathematical and General, 15(10):3241, 1982.

[47] Jascha Sohl-Dickstein, Peter B Battaglino, and Michael R DeWeese. New method for parameter estimation in probabilistic models: minimum probability flow. Physical Review Letters, 107(22):220601, 2011.

[48] David J Thouless, Philip W Anderson, and Robert G Palmer. Solution of ‘solvable model of a spin glass’. Philosophical Magazine, 35(3):593–601, 1977.

[49] Yasser Roudi, Joanna Tyrcha, and John Hertz. Ising model for neural data: model quality and approximate methods for extracting functional connectivity. Physical Review E, 79(5):051915, 2009.

[50] Vitor Sessak and Róemi Monasson. Small-correlation expansions for the inverse Ising problem. Journal of Physics A: Mathematical and Theoretical, 42(5):055001, 2009.

[51] Andrew Y. Ng. Feature selection, l1 vs. l2 regularization, and rotational invariance. In Proceedings of the Twenty-first International Conference on Machine Learning, ICML ’04, pages 78-, New York, NY, USA, 2004. ACM.

[52] Su in Lee, Honglak Lee, Pieter Abbeel, and Andrew Y. Ng. Efficient l1 regularized logistic regression. In In Proceedings of the Twenty-first National Conference on Artificial Intelligence (AAAI-06, pages 1–9, 2006.

[53] Mark Schmidt, Glenn Fung, and Rómer Rosales. Fast optimization methods for L_1_ regularization: A comparative study and two new approaches. In Machine Learning: ECML 2007, pages 286–297. Springer, 2007.

[54] Aurlien Decelle, Federico Ricci-Tersenghi, and Pan Zhang. Data quality for the inverse lsing problem. Journal of Physics A: Mathematical and Theoretical, 49(38):384001, 2016.

[55] Jonathan W Pillow and Peter E Latham. Neural characterization in partially observed populations of spiking neurons. In Advances in Neural Information Processing Systems, pages 1161–1168, 2007.

[56] Duane Q Nykamp. Revealing pairwise coupling in linear-nonlinear networks. SIAM Journal on Applied Mathematics, 65(6):2005–2032, 2005.

[57] Daniel Soudry, Suraj Keshri, Patrick Stinson, Min-hwan Oh, Garud Iyengar, and Liam Paninski. Efficient “shotgun” inference of neural connectivity from highly sub-sampled activity data. PLoS Comput Biol, 11(10):e1004464, 2015.

[58] Jayant E Kulkarni and Liam Paninski. Common-input models for multiple neural spike-train data. Network: Computation in Neural Systems, 18(4):375–407, 2007.

[59] Duane Q Nykamp. A mathematical framework for inferring connectivity in probabilistic neuronal networks. Mathematical Biosciences, 205(2):204–251, 2007.

[60] Joanna Tyrcha and John Hertz. Network inference with hidden units. arXiv preprint arXiv:1301.7274, 2013.

[61] Michael Vidne, Yashar Ahmadian, Jonathon Shlens, Jonathan W Pillow, Jayant Kulkarni, Alan M Litke, EJ Chichilnisky, Eero Simoncelli, and Liam Paninski. Modeling the impact of common noise inputs on the network activity of retinal ganglion cells. Journal of computational neuroscience, 33(1):97–121, 2012.

[62] John Widloski, Michael P Marder, and Ila R Fiete. Inferring circuit mechanisms from sparse neural recording and global perturbation in grid cells. eLife, 7:e33503, 2018.

[63] Dmitriy Aronov, Lena Veit, Jesse H Goldberg, and Michale S Fee. Two distinct modes of forebrain circuit dynamics underlie temporal patterning in the vocalizations of young songbirds. Journal of Neuroscience, 31(45):16353–16368, 2011.

[64] Jose Casadiego, Mor Nitzan, Sarah Hallerberg, and Marc Timme. Model-free inference of direct network interactions from nonlinear collective dynamics. Nature communications, 8(1):2192, 2017.

[65] I. Nemenman, F. Shafee, and W. Bialek. Entropy and inference, revisited. In T. G. Dietterich, S. Becker, and Z. Ghahramani, editors, Advances in Neural Information Processing Systems 14, pages 471–478, Cambridge, MA, 2002. MIT Press.

[66] Andrea Montanari and Jose A Pereira. Which graphical models are difficult to learn? In Advances in Neural Information Processing Systems, pages 1303–1311, 2009.

[67] Federico Ricci-Tersenghi. The Bethe approximation for solving the inverse Ising problem: a comparison with other inference methods. Journal of Statistical Mechanics: Theory and Experiment, 2012(08):P08015, 2012.

[68] Hans A Bethe. Statistical theory of superlattices. In Proc. Roy. Soc. London A, volume 150, pages 552–575, 1935.

[69] Marc Mézard and Giorgio Parisi. The Bethe lattice spin glass revisited. The European Physical Journal B-Condensed Matter and Complex Systems, 20(2):217–233, 2001.

[70] Yasser Roudi, Erik Aurell, and John A Hertz. Statistical physics of pairwise probability models. Frontiers in computational neuroscience, 3, 2009.

[71] Steven P Strong, Roland Koberle, Rob R de Ruyter van Steveninck, and William Bialek. Entropy and information in neural spike trains. Physical Review Letters, 80(1):197, 1998.

[72] Marlene R Cohen and Adam Kohn. Measuring and interpreting neuronal correlations. Nature neuroscience, 14(7):811–819, 2011.

[73] Thierry Mora and William Bialek. Are biological systems poised at criticality? Journal of Statistical Physics, 144(2):268–302, 2011.

[74] Liam Paninski. Estimating entropy on m bins given fewer than m samples. IEEE Transactions on Information Theory, 50(9):2200–2203, 2004.

